# Nuclear corepressors NCOR1 and NCOR2 entrain thymocyte signaling, selection, and emigration

**DOI:** 10.1101/2023.09.27.559810

**Authors:** Natalie A. David, Robin D. Lee, Rebecca S. LaRue, Sookyong Joo, Michael A. Farrar

## Abstract

T cell development proceeds via discrete stages that require both gene induction and gene repression. Transcription factors direct gene repression by associating with corepressor complexes containing chromatin-remodeling enzymes; the corepressors NCOR1 and NCOR2 recruit histone deacetylases to these complexes to silence transcription of target genes. Earlier work identified the importance of NCOR1 in promoting the survival of positively-selected thymocytes. Here, we used flow cytometry and single-cell RNA sequencing to identify a broader role for NCOR1 and NCOR2 in regulating thymocyte development. Using *Cd4-cre* mice, we found that conditional deletion of NCOR2 had no effect on thymocyte development, whereas conditional deletion of NCOR1 had a modest effect. In contrast, *Cd4-cre* x *Ncor1^f/f^* x *Ncor2^f/f^* mice exhibited a significant block in thymocyte development at the DP to SP transition. Combined NCOR1/2 deletion resulted in increased signaling through the T cell receptor, ultimately resulting in elevated BIM expression and increased negative selection. The NF-κB, NUR77, and MAPK signaling pathways were also upregulated in the absence of NCOR1/2, contributing to altered CD4/CD8 lineage commitment, TCR rearrangement, and thymocyte emigration. Taken together, our data identify multiple critical roles for the combined action of NCOR1 and NCOR2 over the course of thymocyte development.

## Introduction

Early T cell progenitors migrate from the bone marrow to the thymus to complete their development. Once in the thymus, they proceed through a sequence of tightly regulated developmental steps and stringent checkpoints. A developing thymocyte must rearrange its TCRβ locus, proliferate, rearrange its TCRα locus, undergo positive selection, commit to the CD4 or CD8 lineage, avoid negative selection, and emigrate from the thymus to successfully enter the periphery as a naïve T cell^1^. These steps ensure a diverse TCR repertoire while limiting the potential for self-reactivity^2^. These steps must be induced sequentially, and each is governed by a distinct transcriptional program. Careful regulation of this process is critical for thymocyte development^3^. While considerable effort has focused on how gene transcription directs T cell development, less is known about how gene repression contributes to appropriate differentiation.

The process of gene repression involves the interaction of transcription factors with corepressor complexes, which direct changes to the epigenetic landscape to silence gene expression^4^. The corepressors NCOR1 and NCOR2 were first identified by their interactions with nuclear receptors^5,6^. Since their initial discovery, they are now recognized to bind a wide variety of transcription factors, including several involved in lymphocyte development such as STAT5, NF-κB, AP-1, and NUR77^7–13^. NCOR1 and NCOR2 serve as a scaffold for the recruitment of histone deacetylases, particularly HDAC3, to DNA-bound transcription factors^14–20^. Beyond enabling assembly of a repressor complex, they are required to activate the deacetylase function of HDAC3^21,22^. The repressor complex silences gene transcription by decreasing chromatin accessibility through histone deacetylation. Here we demonstrate multiple roles for NCOR1 and NCOR2 in silencing gene expression during thymocyte development. Previous studies have established that NCOR1 plays an important role in repressing *Bcl2l11* (BIM) expression and promoting the survival of positively-selected thymocytes^23,24^. Deletion of HDAC3, another member of the NCOR1/2 corepressor complex, results in a broader phenotype. Early deletion of HDAC3 using *Cd2-cre* x *Hdac3^f/f^* mice also increased *Bcl2l11* expression, but this was not observed in *Cd4-Cre x Hdac3^f/f^* mice. In contrast, *Cd2-cre x Hdac3f/f* and *Cd4-Cre x Hdac3^f/f^* mice exhibited a broad range of defects in T cell development and maturation, respectively^25–28^. Since loss of HDAC3 has a more severe phenotype than loss of NCOR1, these findings suggest potential functional redundancy between NCOR1 and NCOR2. To understand how nuclear corepressors NCOR1 and NCOR2 coordinate T cell development, we generated *Cd4-cre* x *Ncor1^f/f^*, *Cd4-cre* x *Ncor2^f/f^*, and *Cd4-cre* x *Ncor1^f/f^*x *Ncor2^f/f^* mice. Our work identified key roles for NCOR1 and NCOR2 in shaping thymocyte survival, selection, TCR repertoire, and emigration.

## Results

Germline deletion of NCOR1 or NCOR2 is embryonic lethal^29–31^. To evaluate the impact of NCOR1 and NCOR2 deletion in T cells, we utilized a conditional knockout system with existing *Ncor1^f/f^* and *Ncor2^f/f^* mice^24,32^. We generated *Cd4-cre* x *Ncor1^f/f^*, *Cd4-cre x Ncor2^f/f^*, *Cd4-cre* x *Ncor1^f/f^* x *Ncor2^f/f^*, and *Cd4-creER^T^*^2^ x *Ncor1^f/f^* x *Ncor2*^f/f^ mice for this study. The NCOR-deficient mice did not demonstrate increased morbidity or mortality in specific-pathogen free (SPF) conditions and exhibited comparable growth to their wild-type (WT) littermates (data not shown). NCOR1 deletion was confirmed by flow cytometry (Supplemental Figure 1A), though we were not able to do the same for NCOR2 due to lack of specific anti-NCOR2 flow cytometry antibodies. Due to residual NCOR1 protein expression in the CD4+ CD8+ double positive (DP) thymocyte population (primarily from TCRβ-DPs), we elected to pre-gate on TCRβ+ cells for all subsequent analyses unless otherwise specified (Supplemental Figure 1B). *Cd4-cre x Ncor2^f/f^* mice did not exhibit any obvious phenotype and were not further investigated (data not shown). Numbers of CD4 and CD8 SP thymocytes trended lower in *Cd4-cre x Ncor1^f/f^* (NCOR1 KO) mice but did not reach statistical significance. In contrast, NCOR1 and NCOR2 conditional double knockout (DKO) mice demonstrated a significant block in thymocyte development (Figure 1A). There was a marked increase in the frequency of DP cells and decrease in the frequency and number of both CD4 single positive (SP) and CD8 SP cells in the absence of both NCORs (Figure 1B). CD4 SP cells were more severely impacted than CD8 SP cells (8-fold vs 3.6-fold reduction in DKO mice), resulting in a decreased CD4:CD8 ratio (Figure 1C). Consistent with defective T cell development in the thymus, we also observed a dramatic decline in the CD4+ and CD8+ T cell populations in the spleens of DKO mice (Supplemental Figure 2A). As in the thymus, splenic CD4+ T cells were more severely impacted by the loss of NCOR1/2 than CD8+ T cells (17.5-fold reduction vs 4-fold reduction), resulting in a decreased CD4:CD8 ratio (Supplemental Figure 2B).

**Figure 1.**
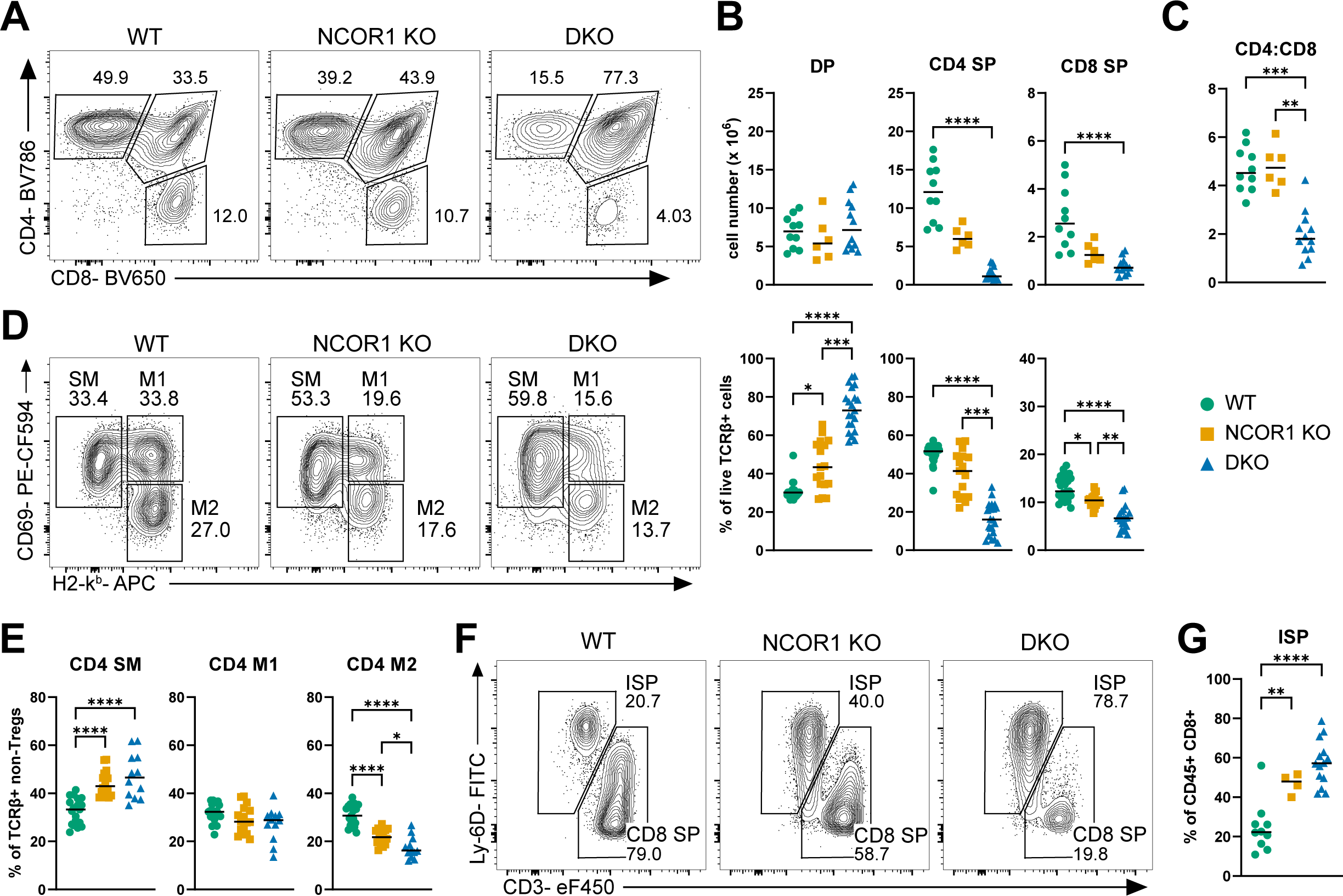
Corepressors NCOR1 and NCOR2 are required for normal T cell development and maturation. (A) Representative flow cytometry plots of CD4 single positive (CD4 SP), CD8 single positive (CD8 SP), and CD4 CD8 double positive (DP) thymocytes isolated from WT, *Cd4-cre x Ncor1^f/f^* (NCOR1 KO), and *Cd4-cre x Ncor1^f/f^ x Ncor2^f/f^* (DKO). Cells were pre-gated on live TCRβ+. (B) Summary data of CD4 SP, CD8 SP, and DP population counts (top panel) and frequencies (bottom panel). (C) Ratio of CD4 SP to CD8 SP thymocytes. (D) Representative flow cytometry plots of CD4 SP intermediate populations: semi-mature (SM), mature 1 (M1), and mature 2 (M2), distinguished by H2-k^b^ (MHC class I) and CD69 expression. Pre-gated on live TCRβ+ CD4+ CD8-CD73-CD25-Foxp3-cells. (E) Summary data for frequencies of CD4 SM, M1, and M2 intermediates. (F) Representative flow cytometry plots of CD8 SP and immature SP (ISP) thymocytes, identified by CD3 and Ly-6D expression, from WT, NCOR1 KO, and DKO mice. Pre-gated on live CD45+ CD4-CD8+ cells. (G) Summary data for frequency of ISP population. Data were compiled from a minimum of 4 independent experiments for all panels. Horizontal bars represent the median. Statistical significance was determined by Kruskal Wallis test with Dunn’s multiple comparison test (B, C) or one-way ANOVA with Tukey’s multiple comparison test (E, G). *p < 0.05, **p < 0.01, ***p < 0.001, ****p < 0.0001.

To more precisely identify the timing of the thymic developmental block, we examined intermediate CD4 SP populations^33^. After excluding TCRβ-cells, CD73+ mature recirculating cells, and regulatory T cells, we used MHC-I (H2-k^b^) and CD69 expression to subset CD4 SP thymocytes into three subpopulations: CD69^+^MHC-I^−^ (semi-mature; SM), CD69^+^MHC-I^+^ (mature 1; M1), and CD69^−^MHC-I^+^ (mature 2; M2) cells (Figure 1D). NCOR1 KO and DKO mice demonstrated a significant increase in the frequency of CD4 SP SM cells and a decrease in the frequency of CD4 SP M2 cells (Figure 1E), although the defect in DKO mice was more pronounced. Finally, we used CD3 and Ly-6D to distinguish immature single positive (ISP) cells from CD8 SP cells in CD45+ CD8+ thymocytes (Figure 1F)^34^. The loss of NCOR1 and NCOR2 resulted in a substantial increase in the frequency of ISP thymocytes relative to CD8 SP thymocytes, consistent with a defect in CD8 SP development (Figure 1G). These data demonstrate that loss of NCOR1 and NCOR2 significantly impairs T cell development and leads to a dramatic reduction in both CD4 SP and CD8 SP thymocytes.

Following our initial characterization of WT and NCOR-deficient thymocytes by flow cytometry, we performed a single cell RNA-seq (scRNA-seq) experiment to understand which molecular processes are impacted by the loss of corepressors NCOR1 and NCOR2 (Figure 2A). We harvested thymi from two (1 male, 1 female) 6-week-old mice from each genotype (WT, NCOR1 KO, and DKO). Thymocytes from each mouse were hashtagged with a unique barcode. We sorted TCRβ+ CD4+ CD8+ cells (DP), TCRβ+ CD4+ CD73-CD25-GITR-cells (CD4 SP), and TCRβ+ CD4+ CD73-cells that are CD25+ and/or GITR+ (regulatory T cell progenitors and regulatory T cells). These populations were combined in equal proportion immediately prior to sequencing. As a result, CD4 SP thymocytes were enriched relative to DP thymocytes. Using the 10X Genomics platform, we generated 5’ scRNA-seq libraries. This allowed us to collect concurrent transcriptome and TCR sequencing data. After removing multiplets, a total of 11,216 cells were captured in this experiment. All genotypes were well-represented in the population of captured cells: 4187 WT, 3716 NCOR1 KO, and 3313 DKO (Figure 2B). We compared multiple resolutions for unsupervised clustering in Seurat and selected resolution 1.4 for analysis (Supplemental Figure 3A). Genes with variable patterns of expression across T cell development were used to assign cluster identities (Supplemental Figure 3B, Supplemental Figure 4A). Although we enriched for regulatory T cells and regulatory T cell progenitors in our experimental design, analysis of these populations is beyond the scope of this publication.

**Figure 2.**
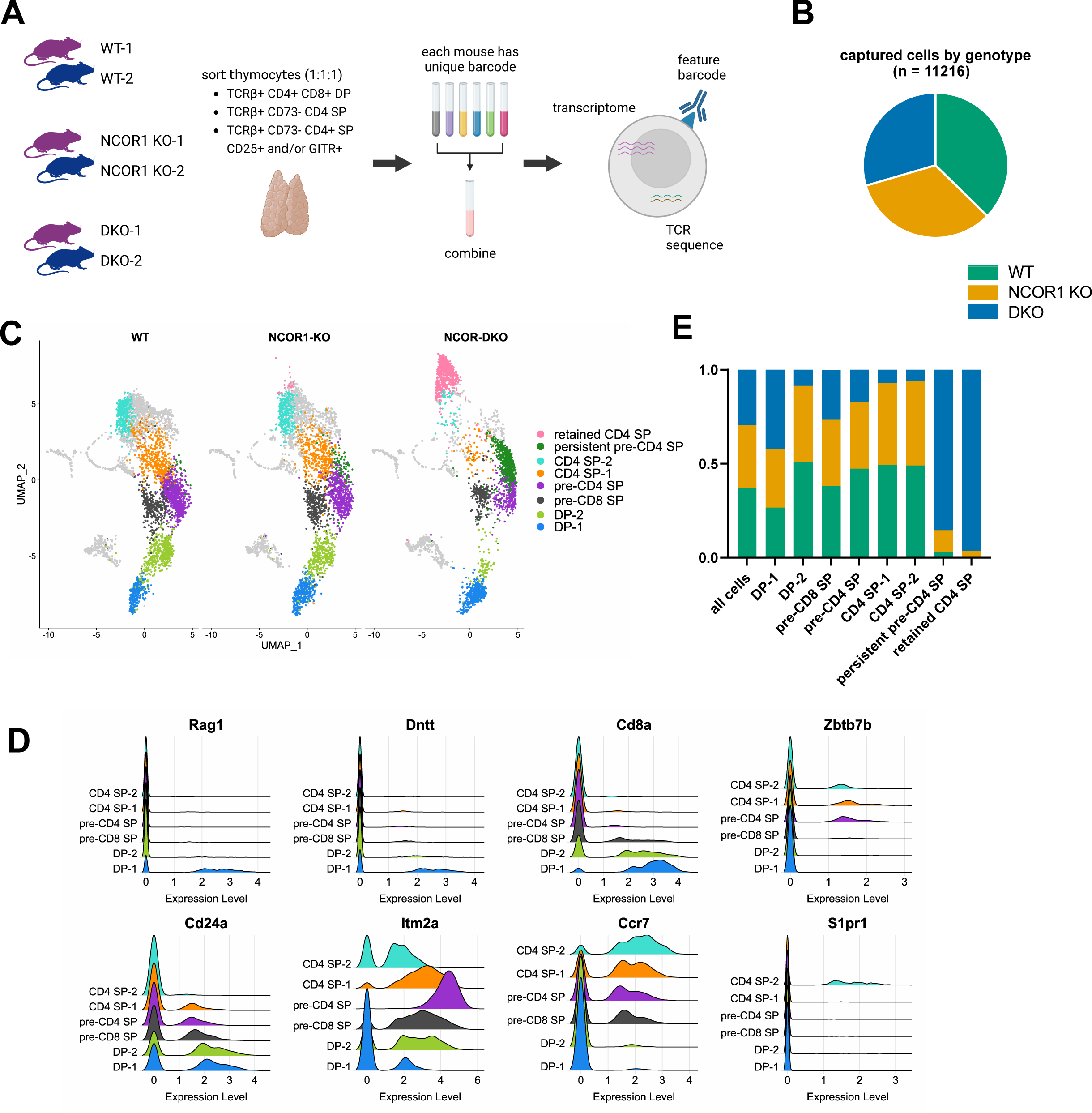
scRNA-seq reveals altered composition and distribution of DP and CD4 SP thymocyte populations in absence of NCOR1/2. (A) Schematic of scRNA-seq experiment, including TCR sequencing and feature barcoding, using two 6-week-old mice per genotype. One male and one female were used for each genotype. (B) Total number of cells captured after doublet removal and proportion of total population derived from each genotype. Each genotype label is composed of two biological replicates as detailed in A. (C) UMAP plot of scRNAs-seq data, split by genotype, highlighting only the clusters that were investigated in this study. (D) Ridge plot for select genes whose expression changes over the course of T cell development and were used to assign cluster identities. (E) Contribution of cells from each genotype within specified clusters.

We focused our study on clusters related to the DP to SP transition, as well as the selection and maturation of CD4 SP thymocytes (Figure 2C). The selected clustering resolution divided *Cd4+ Cd8+* expressing cells into two populations, which we termed DP-1 and DP-2. *Cd4+ Cd8-* cells with low expression of maturation markers fell into two selection intermediate populations, pre-CD4 SP and pre-CD8 SP, that are CD4- and CD8-fated, respectively. *Cd4+ Cd8-* cells with increasing expression of maturation markers were divided into CD4 SP-1 and CD4 SP-2. These assignments were supported by well-established patterns of gene expression across T cell development (Figure 2D). For example, *Rag1* and *Dntt* were expressed in the least mature DP population, DP-1. *Cd8a* expression decreased from DP-1 to DP-2 to pre-CD8 SP clusters, and was almost completely extinguished in pre-CD4 SP, CD4 SP-1, and CD4 SP-2 clusters. *Zbtb7b* (which encodes THPOK) expression indicates commitment to the CD4 SP lineage and was expressed only in the pre-CD4 SP, CD4 SP-1, and CD4 SP-2 subsets. *Cd24a* is continuously downregulated as cells mature^35^; this pattern was clearly observed in our dataset as well. *Itm2a* expression is induced during thymocyte selection^36^; consistent with those findings, we observed increased *Itm2a* expression in the pre-CD8 SP and pre-CD4 SP populations relative to the DP-1 and DP-2 populations. Increasing *Ccr7* expression is consistent with increasing thymocyte maturity while induction of *S1pr1* expression indicates the most mature CD4 SP cells^37,38^; this pattern was also observed in our scRNA-seq clusters. DKO cells were severely underrepresented in the DP-2, CD4 SP-1, and CD4 SP-2 clusters and more moderately underrepresented in the pre-CD8 SP and pre-CD4 SP clusters (Figure 2E). The more moderate reduction in pre-CD4 SP and pre-CD8 SP subsets likely reflects the overall less mature phenotype of NCOR-deficient CD4 SPs, as identified in Figure 1. Finally, two clusters were comprised almost exclusively of DKO cells (Figure 2E, Supplemental Figure 4A). The “persistent pre-CD4 SP” subset was most similar to the pre-CD4 SP subset but exhibited ongoing, robust TCR signaling as indicated by elevated *Cd5, Nr4a1*, and *Dusp5* expression (Supplemental Figure 4B). The second DKO predominant cluster, which we called the “retained CD4 SP” cluster, had a very distinctive phenotype and is addressed in Figure 6.

While multiple aspects of thymocyte development were impaired in NCOR-deficient mice, the defect in the DP to SP transition was particularly pronounced. Signaling through a rearranged TCR is required for a thymocyte to advance beyond the DP stage and commit to the CD4 or CD8 lineage. We explored whether impaired signaling was contributing to the observed defect in positive selection in DKO mice. Gene set enrichment analysis (GSEA) of the DP-1 cluster identified an increase in genes associated with T cell activation in the conditional NCOR1 KO and DKO mice (Figure 3A). We also discovered significant enrichment of pathways induced by TCR signaling, including NF-κB and MAPK in NCOR-deficient thymocytes in the DP-1, DP-2, pre-CD SP, and CD4 SP-1 clusters (Supplemental Figure 5A, B). These data implicate NCOR1/2 in repressing signaling through the NF-κB and MAPK pathways.

**Figure 3.**
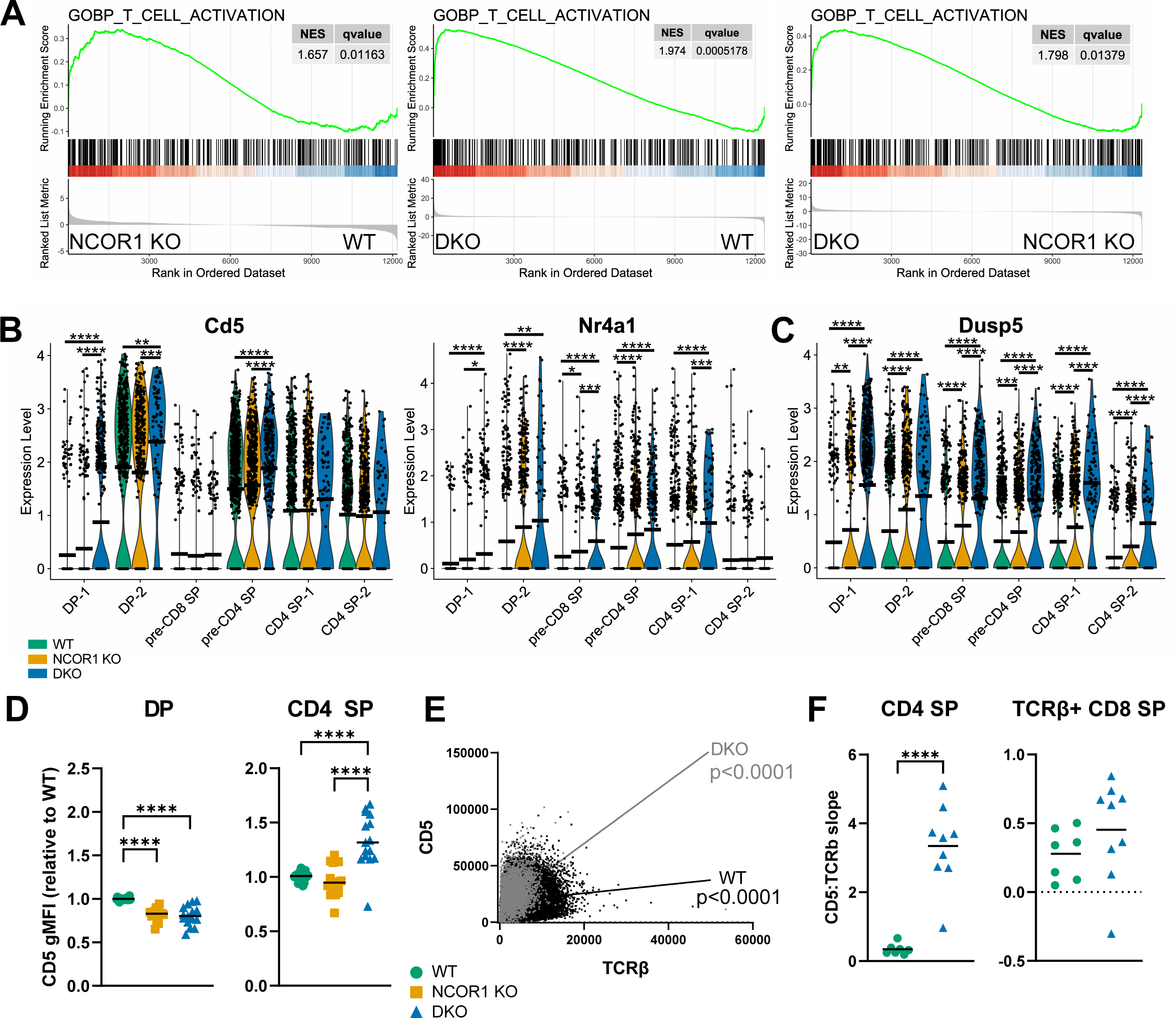
CD4 SP thymocytes demonstrate enhanced signaling through their T cell receptors in the absence of NCOR1 and NCOR2. (A) Enrichment for GOBP T cell activation gene signature in DP-1. GSEA was completed using differential expression lists between genotypes within this cluster: WT and NCOR1 KO (left), WT and DKO (center), and NCOR1 KO and DKO genotypes (right). NES = normalized enrichment score. (B) Violin plots of *Cd5* (left) and *Nr4a1* (NUR77; right) expression across specified scRNA-seq clusters. (C) Violin plot of *Dusp5* expression across specified scRNA-seq clusters. (D) Geometric mean fluorescence intensity (gMFI) of CD5 in TCRβ+ DP and TCRβ+ CD4 SP thymocytes by flow cytometry. Values were normalized to WT sample(s) within each experiment. (E) Correlation of CD5 and TCRβ scale values in CD4 SP thymocytes. Each dot represents a single cell from one WT (black) or DKO (gray) sample. P value indicates whether the slope for linear regression is significantly non-zero. (F) Summary data of linear regression slopes between CD5 and TCRβ in CD4 SP (left) and CD8 SP (right) thymocytes from WT and DKO mice. CD4 SP thymocytes were not pre-gated on TCRβ+. CD8 SP thymocytes were pre-gated on TCRβ+. Flow cytometry data were compiled from 5 independent experiments. Horizontal bars represent the mean (B, C) or median (D). Statistical significance was determined by two-tailed Wilcoxon rank-sum test (B, C), one-way ANOVA with Tukey’s multiple comparison test (D), or unpaired, two-tailed t test (F). *p < 0.05, **p < 0.01, ***p < 0.001, ****p < 0.0001.

Our GSEA data suggest that TCR signaling is enhanced in the absence of NCOR1/2. Consistent with this idea, we found that *Cd5* and *Nr4a1*, genes whose expression has been shown to correlate with TCR signal strength, were significantly upregulated across DP and CD4 SP clusters in NCOR1 KO and DKO mice relative to WT controls^39–42^ (Figure 3B). The expression of *Dusp5*, another gene robustly induced by TCR signaling, was elevated throughout development in NCOR-deficient mice as well (Figure 3C)^43^. The finding of *Cd5* upregulation was validated using flow cytometry, as CD5 expression was also significantly increased on TCRβ+ CD4 SP cells in the absence of NCOR1/2 (Figure 3D). Finally, CD5 surface expression has been shown to be proportional to both surface level TCR expression and TCR signaling intensity^39^. We performed linear regression to relate TCRβ and CD5 protein expression on all CD4 SP thymocytes. There was a significantly higher level of CD5 expression relative to TCRβ expression in the DKO mice (Figure 3E, F). In addition to the elevated CD5 expression in the absence of NCORs, TCRβ expression was significantly reduced in CD4 SP thymocytes (Supplementary Figure 5C, D). This relationship between TCRβ and CD5 expression was not observed in TCRβ+ CD8 SP thymocytes from DKO mice (Figure 3F). Our results suggest that NCOR1 and NCOR2 are required for attenuation of TCR signaling specifically during CD4 SP thymocyte development.

Previous studies identified an important role for NCOR1 in promoting thymocyte survival^23,24^. We also found increased apoptosis in the absence of NCOR1 and NCOR2. GSEA of the DP-1 population identified significant enrichment for an apoptotic gene signature in DKO relative to WT, and in DKO relative to NCOR1 KO (Figure 4A). Consistent with earlier work in which deletion of NCOR1 resulted in de-repression of the pro-apoptotic gene *Bcl2l11* (BIM), we observed a significant increase in *Bcl2l11* expression in NCOR1 KO thymocytes, and a further significant increase in DKO thymocytes relative to NCOR1 KO alone^23^ (Figure 4B). Elevated *Bcl2l11* expression in the absence of NCOR1 and NCOR2 was particularly pronounced in the DP-1 and DP-2 clusters, though it persisted through the CD4 SP-1 stage. These observations were validated at a protein level using flow cytometry. BIM expression was elevated in TCRβ+ DP and across all TCRβ+ CD4 SP thymocyte subsets in DKO mice (Figure 4C). These results illustrate that BIM is induced to higher levels in the absence of NCORs, beginning with DP thymocytes and continuing through CD4 SP maturation. Possibly to compensate for their tenuous survival, TCRβ+ CD4 thymocytes from NCOR1 KO mice have also been shown to express higher levels of the anti-apoptotic protein BCL2^24^. We observed a comparable rise in *Bcl2* expression in the pre-CD8 SP and pre-CD4 SP clusters (Figure 4D). Increasing *Bcl2* expression may rescue a small population of NCOR-deficient cells, allowing them to survive and advance beyond the DP stage.

**Figure 4.**
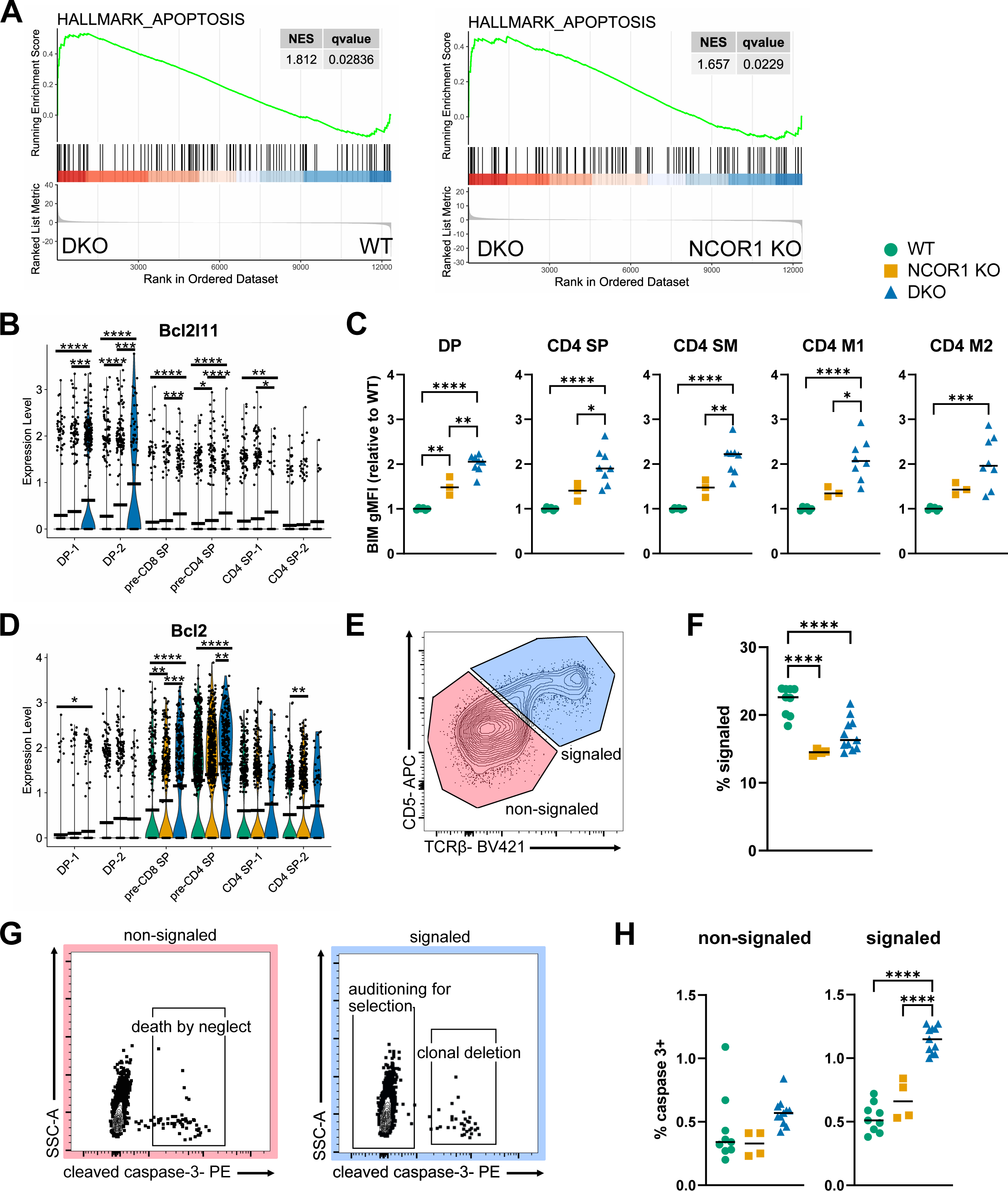
Apoptosis and negative selection are enhanced in the absence of NCOR1 and NCOR2. (A) Enrichment for hallmark apoptosis gene signature. GSEA was performed using differential expression lists between WT and DKO (left) and NCOR1 KO and DKO (right) samples within the DP-1 cluster. NES = normalized enrichment score. (B) Violin plots of *Bcl2l11* (BIM) expression across select clusters from scRNA-seq dataset. (C) BIM gMFI by flow cytometry in TCRβ+ DP (left), all TCRβ+ CD4 (second from left), CD4 SM (center), CD4 M1 (second from right), and CD4 M2 (right). Values were normalized to WT sample(s) within each experiment. CD4 SP SM, M1, and M2 were gated as shown in Figure 1D. (D) Violin plot of *Bcl2* expression across select clusters from scRNA-seq dataset. (E) Representative flow cytometry plot distinguishing signaled (CD5+ TCRβ+) and non-signaled (CD5-TCRβ-) thymocytes. Pre-gated on live single cells. (F) Summary data for frequency of signaled thymocytes across genotypes. (G) Representative flow cytometry plots of cleaved caspase-3+ staining for populations gated in (E). Cleaved caspase-3+ non-signaled cells identified thymocytes undergoing death by neglect (left) and cleaved caspase-3+ signaled cells identified thymocytes undergoing clonal deletion (right). (H) Summary data for frequency of cleaved caspase-3+ cells among non-signaled (left) and signaled thymocytes (right). Each symbol represents an individual mouse. Flow cytometry data were compiled from a minimum of 5 independent experiments. Horizontal bars indicate mean (B, D) or median (C, F, H). Statistical significance was determined by two-tailed Wilcoxon rank-sum test (B, D) or one-way ANOVA with Tukey’s multiple comparison test (C, F, H). *p < 0.05, **p < 0.01, ***p < 0.001, ****p < 0.0001.

Cell death and TCR signaling are closely related in the thymus. The strength of the self-peptide:MHC and TCR interaction determines the fate of a developing thymocyte: weak TCR signaling results in death by neglect, intermediate TCR signaling results in positive selection, and strong TCR signaling results in clonal deletion. Based on our observations of both significantly increased TCR signaling and increased apoptosis in the absence of NCOR1/2, we used a clonal deletion assay to measure negative selection in the thymus^44^. TCRβ and CD5 co-expression were used to identify signaled thymocytes (Figure 4E). All other cells were considered non-signaled. There was a modest reduction in the frequency of signaled cells in NCOR-deficient thymocytes (Figure 4F). Of the signaled thymocytes, cleaved caspase-3+ cells are undergoing clonal deletion, while cleaved caspase-3+ non-signaled thymocytes are undergoing death by neglect (Figure 4G). There was a striking increase in the frequency of cleaved caspase-3+ signaled cells in DKO thymocytes relative to their WT and NCOR1 KO counterparts (Figure 4H). These results indicate that enhanced negative selection, likely secondary to amplified TCR signaling, drives increased cell death in the absence of NCOR1 and NCOR2.

Having identified changes in TCR signaling, we next examined our single-cell TCR-seq dataset to investigate whether NCOR deficiency was associated with an altered TCR repertoire. Generating a diverse TCR repertoire is an iterative process involving multiple rounds of somatic gene rearrangement. The β-chain locus rearranges first, during the CD4-CD8-double negative (DN) stage. The process begins with D_β_ to J_β_ rearrangements followed by V_β_ to DJ_β_ rearrangements (Figure 5A). Rearrangements continue, using increasingly distal gene segments, until a productive (i.e., in-frame) β-chain is generated^45^. During the DP stage, the α-chain locus undergoes rearrangement in a parallel process with its V_α_ to J_α_ gene segments (Figure 5B). Secondary V_α_ to J_α_ gene rearrangements occur until the thymocyte is positively or negatively selected, or dies^46^; generation of an in-frame α-chain is insufficient to stop the process. In our conditional knockout mice, TCRβ rearrangement takes place in the double negative (DN) stage, prior to NCOR deletion, and TCRα rearrangement begins shortly after deletion is induced at the DP stage. If loss of NCOR1 and NCOR2 had an impact on the TCR repertoire, it should only be reflected at the TCRα locus.

**Figure 5.**
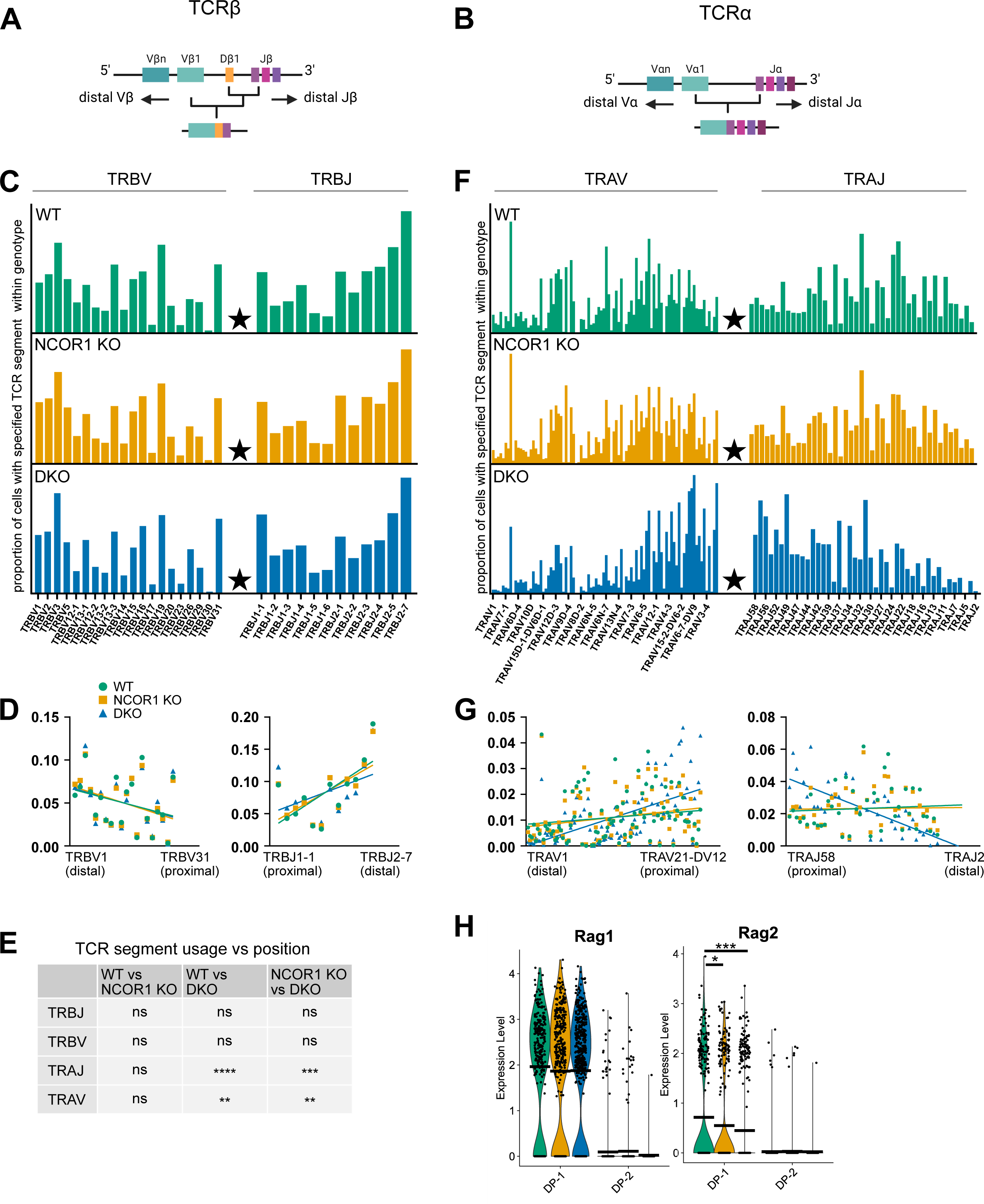
Loss of NCOR1 and NCOR2 results in a proximally-biased TCRα repertoire. (A) Schematic of TCRβ locus demonstrating initial D to J recombination and V to DJ recombination and direction of subsequent recombination events. (B) Schematic of TCRα locus illustrating initial V to J recombination and direction of subsequent recombination events. (C) Distribution of segment utilization in WT (top), NCOR1 KO (middle), and DKO (bottom) thymocytes for TCRβ locus. Star indicates initial recombination site. TCR segments are organized 5’ to 3’. Bar height indicates the proportion of cells within each genotype with a given TCR segment indicated on the x-axis. (D) Linear regression of TRBV (left) and TRBJ (right) segment position, relative to initial recombination site, and proportion of cells with this segment found in each genotype. (E) Comparison of relationship between distribution of TCR segment utilization and position (slope from linear regression) between genotypes. (F) Same as C using data from TCRα locus. (G) Linear regression of TRAV (left) and TRAJ (right) segment position, relative to initial recombination site, and proportion of cells with this segment found in each genotype. (H) Violin plot of *Rag1* (left) and *Rag2* (right) expression in DP-1 and DP-2 thymocytes. Statistical significance was determined by one-way ANOVA with Tukey’s multiple comparison test (E) or two-tailed Wilcoxon rank-sum test (H). Horizonal bars indicate mean (H). *p < 0.05, **p < 0.01, ***p < 0.001, ****p < 0.0001.

We began with the TCRβ locus by comparing the proportion of cells with a given V_β_ or J_β_ gene segment among all cells of each genotype (Figure 5C). D_β_ segments were excluded from our analysis due to their limited diversity. All of the gene segments were ordered 5’ to 3’ and assigned a distance for each gene segment relative to the site of the initial V_β_ to DJ_β_ rearrangement. We performed a linear regression to relate the frequency with which a segment is used to its proximity to the initial rearrangement site (Figure 5D). The TRBV and TRBJ slopes were not significantly different between genotypes (Figure 5E). This was the expected result as NCOR1 and NCOR2 are still present during TCRβ rearrangement in DN thymocytes. We then moved to the TCRα locus and compared the distribution of V_α_ and J_α_ gene segment utilization within each genotype (Figure 5F). Linear regression was again completed to relate gene segment utilization and distance from the initial V_α_ to J_α_ rearrangement (Figure 5G). We observed a strong bias toward proximal TRAV and TRAJ gene segments in the DKO repertoire (Figure 5E). There was no relationship between TCRα gene segment usage and distance in the WT and NCOR1 KO repertoires. Thus, deletion of both NCOR1 and NCOR2 results in selective defects in rearrangement of distal V_α_ and J_α_ genes.

TCRα rearrangement is only stopped by positive selection or apoptosis. The earlier one of these events occurs, the fewer rounds of rearrangement take place, thereby preventing thymocytes from testing more distal V and J segments^46^. Given the increased apoptosis observed in NCOR-deficient thymocytes, we considered the possibility that cells were dying in the DP stage before they had undergone many secondary TCRα rearrangements. However, after examining the TCR repertoire data in the individual clusters, we found no difference in the TRAV or TRAJ slopes between genotypes in the DP-1 and DP-2 populations (Supplemental Figure 6A-C).

*Rag1* expression was not significantly different between genotypes in the DP-1 and DP-2 clusters, though *Rag2* expression was reduced in NCOR-deficient cells in the DP-1 cluster (Figure 5H). Because TCR signaling induces the downregulation of *Rag1* and *Rag2,* the reduced *Rag2* expression observed in this population likely reflects early and robust TCR signaling in the absence of NCORs^47^. Our data support the conclusion that NCOR-deficient thymocytes are fully capable of TCRα rearrangement but are selected after fewer rounds of TCRα rearrangement.

Having compared WT and NCOR-deficient thymocytes across development, we sought to characterize the cluster in our scRNA-seq data that was comprised almost exclusively of DKO cells and lacked WT cells entirely (Supplemental Figure 7A). The top differentially expressed genes in this cluster, relative to all clusters, were *Ly6a* and *Ly6c1* (Supplemental Figure 7B). These genes are not known to be involved in thymocyte development; they are typically used to identify effector and memory T cells^48^. Using flow cytometry, we found an increased frequency of Sca-1+ (Ly-6A/E) and Ly-6C+ CD4 SP thymocytes, particularly in the more mature CD4 SP subsets (Supplemental Figure 7C). These findings suggested that the DKO-enriched cluster was composed of a relatively mature CD4 population. Consistent with maturity, cells in this cluster had *H2-K1, H2-D1,* and *Sell* (CD62L) expression levels at or above those found in the CD4 SP-2 cluster (Supplemental Figure 7D, E). We pondered whether these NCOR-deficient mature T cells had emigrated from the thymus and returned, or whether they had never left the thymus. We had gated out CD73+ T cells prior to sequencing to exclude recirculating T cells and therefore favored a retained CD4 SP hypothesis^49^.

After successfully navigating TCR recombination, positive selection, and negative selection, mature CD4 SP thymocytes must upregulate *S1pr1* and *Klf2* to traffic out of the thymus and into the periphery^38,50,51^. These genes were not only highly expressed in the CD4 SP-2 cluster but also in the DKO-enriched cluster (Figure 6A). This population demonstrated substantial enrichment in a gene set associated with lymphocyte migration as well (Figure 6B). With a transcriptional profile suggesting a capacity to leave, why were these NCOR-deficient mature T cells persisting in the thymus? Downregulation of *Cd69* in mature SP thymocytes is required as it negatively regulates lymphocyte egress^52,53^. Loss of corepressors, including NCOR1 and NCOR2, leads to de-repression of target genes. We identified very high levels of *Cd69* transcript in the DKO-enriched cluster (Figure 6C). CD69 and S1PR1 expression have generally been considered to be mutually exclusive so observing *Cd69* and *S1pr1* co-expression was unexpected^54–56^. At the protein level, CD69 interacts directly with S1PR1 to facilitate internalization and degradation of the receptor^57^. However, using flow cytometry, we noted that S1PR1 and CD69 were simultaneously present on the cell surface of DKO CD4 SP thymocytes (Figure 6D). In the absence of NCOR1 and NCOR2, S1PR1+ CD4 SP thymocytes displayed significantly higher levels of CD69 expression (Figure 6E).

**Figure 6.**
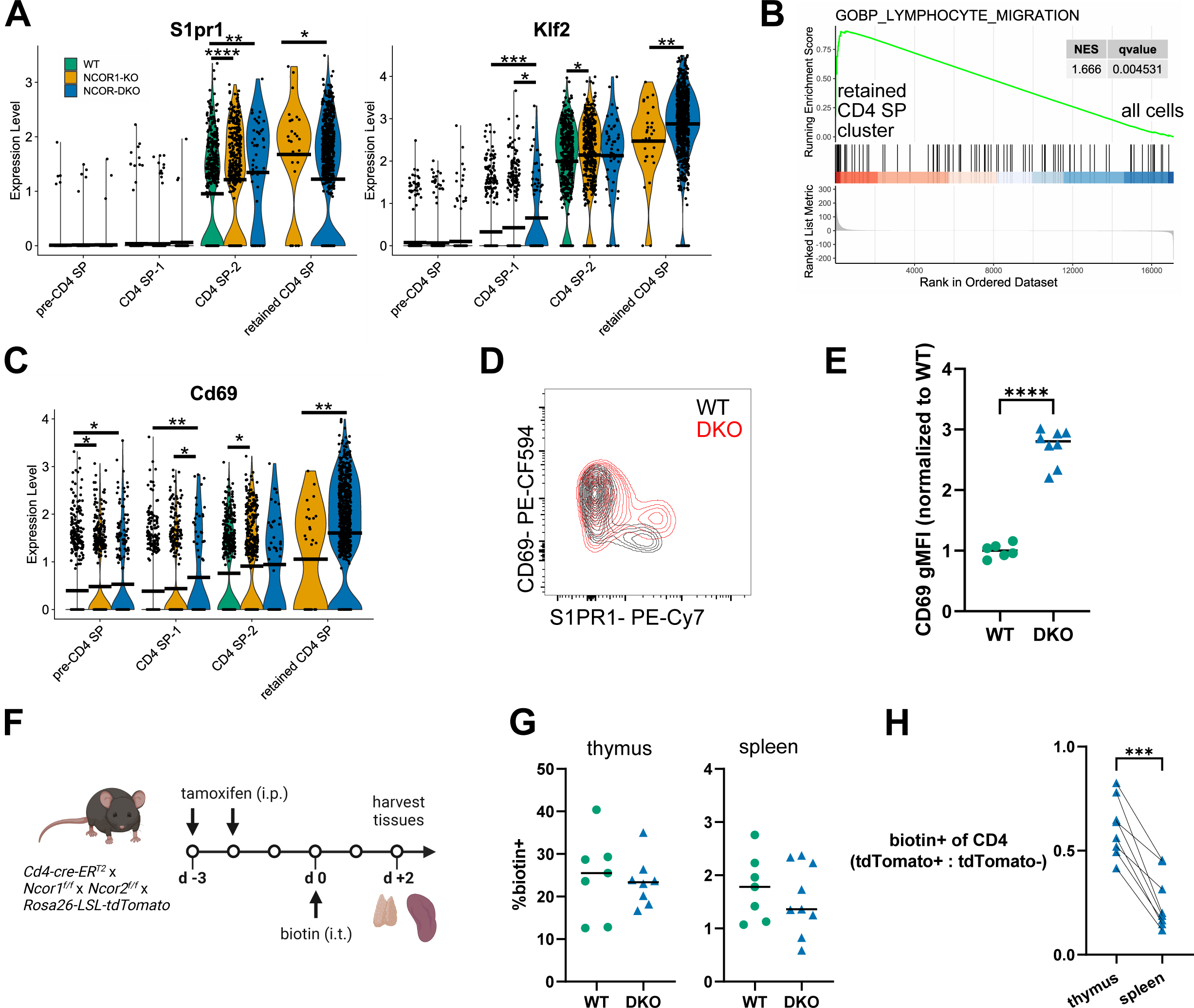
CD4 SP thymocytes upregulate genes required for thymic egress but fail to downregulate CD69 and are retained in the thymus in the absence of NCOR1 and NCOR2. (A) Violin plot of *S1pr1* (left) and *Klf2* (right) expression in pre-CD4 SP and more mature CD4 SP-1 and CD4 SP-2 thymocytes, as well as the cluster comprised largely of DKO cells (“retained CD4 SP”). (B) Enrichment for GOBP lymphocyte migration gene signature. GSEA was completed using a differential expression list between the retained CD4 SP cluster and all other cells. (C) Violin plot of *Cd69* expression in specified clusters from scRNA-seq dataset. (D) Representative flow cytometry plot of CD69 and S1PR1 expression in WT (black) and DKO (red) TCRβ+ CD4 SP thymocytes. (E) Compiled gMFI data for CD69 in S1PR1+ TCRβ+ CD4 SP thymocytes. Values normalized to WT sample(s) within each experiment. (F) Schematic for thymocyte emigration experiment. (G) Frequency of biotin+ TCRβ+ CD4+ CD73-cells in the thymus (left) and biotin+ TCRβ+ CD4+ cells in the spleen (right). (H) Ratio of % biotin+ of tdTomato+ CD4 cells to % biotin+ of tdTomato-CD4 cells. Pre-gated on TCRβ+ CD4+ CD73-(thymus) and TCRβ+ CD4+ (spleen). Flow cytometry and biotin labeling data were compiled from 3 independent experiments. Statistical significance was determined by two-tailed Wilcoxon rank-sum test (A, C), one-way ANOVA with Tukey’s multiple comparison test (E, G), or two-tailed paired t-test (H). Horizonal bars indicate mean (A, C) or median (E, G). *p < 0.05, **p < 0.01, ***p < 0.001, ****p < 0.0001.

To determine whether NCOR1/2-deficient thymocytes have an emigration defect, we developed a strategy to test for impaired thymocyte emigration *in vivo* (Figure 6F). We administered tamoxifen to *Cd4-creER^T^*^2^ *x Ncor1^f/f^ x Ncor2^f/f^ x LSL-tdTomato* mice to induce deletion of NCOR1 and NCOR2. The tdTomato reporter identified cells that successfully deleted NCOR1/2. This treatment regimen resulted in deletion of both NCORs in most, but not all, CD4 SP thymocytes. Next, we performed intra-thymic biotin injection under ultrasound guidance to label thymocytes^58,59^. The mice were sacrificed two days post injection and analyzed by flow cytometry. The frequencies of biotin-labeled TCRβ+ CD4 cells in the thymus (CD4 SP) and spleen (recent thymic emigrants) were not significantly different between WT and DKO mice (Figure 6G). We then looked at differences between biotinylated NCOR-sufficient (tdTomato-) and NCOR-deleted (tdTomato+) cells. In the thymus, approximately 60% of the tdTomato+ cells were biotin+ relative to the tdTomato-cells. In the spleen, only 25% of the tdTomato+ cells were biotin+ relative to the tdTomato-cells (Figure 6H). The significantly lower proportion of biotinylated, NCOR-deficient cells in the spleen compared to the thymus indicated reduced thymic emigration in the absence of these corepressors, as was suggested by our scRNA-seq dataset. Taken together, these results reveal a critical role for NCORs in allowing thymocyte emigration.

## Discussion

Using a conditional knockout model, this study identified novel roles for combined NCOR1 and NCOR2 function in CD4 T cell development. Our study establishes multiple roles for corepressors NCOR1 and NCOR2 in thymocyte differentiation. The developmental perturbations observed in NCOR-deficient thymocytes can be attributed to a confluence of excess activation of multiple signaling pathways, including those involving NF-κB, NUR77, and MAPK. De-repression of these pathways was associated with alterations in thymocyte survival, selection, CD4/CD8 lineage choice, and thymic egress, as well as TCR repertoire and signaling through the T cell receptor.

Previous studies have examined the role of NCOR1 in developing thymocytes. NCOR1 has been shown to bind the *Bcl2l11* promoter and repress *Bcl2l11* transcription, thereby protecting positively-selected thymocytes from negative selection^23^. We too observed increased *Bcl2l11* mRNA and BIM protein expression in DP thymocytes from NCOR 1 KO mice. Although we did not measure *Bcl2l11* expression in thymocytes from NCOR2 KO mice, the overall absence of a phenotype suggests that apoptosis was unaffected by the loss of NCOR2. Considering the extensive interactions between NCORs and transcription factors identified in other biological contexts, it seems unlikely that suppression of apoptosis is the sole role for these proteins in T cells. Consistent with this hypothesis, we observed significantly more perturbation in T cell development in combined NCOR1 and NCOR2 conditional knockout mice than in those with conditional knockout of NCOR1 or NCOR2 alone. For instance, *Bcl2l11* expression was further elevated in DP and CD4 SP thymocytes from DKO mice compared to the increase observed in NCOR1 KO thymocytes. Another finding unique to DKO thymocytes was significantly increased TCR signaling and clonal deletion. TCR stimulation induces early expression of NUR77, proportional to the strength of the TCR signal^42^. NUR77 induces apoptosis in T cells after TCR engagement, partially mediated by its induction of BIM, and is required for efficient clonal deletion^60–65^. NCOR2 is recruited directly to NUR77, so when NCOR2 is deleted, NUR77 activity is de-repressed^12^. The enhanced NUR77 function in the absence of NCOR1/2 presumably drives both higher TCR signaling and increased BIM expression leading to more apoptosis in DKO thymocytes relative to their WT and NCOR1 KO counterparts. Our findings are consistent with increased negative selection in the absence of NCOR1 and NCOR2, possibly via a failure of the NCOR1/2 corepressor complex to limit NUR77-induced BIM-dependent apoptosis.

NCOR1/2 DKO mice also exhibit defects in other aspects of thymocyte development that have not been described in NCOR1 KO mice. For example, we found that NCOR1 and NCOR2 affect CD4/CD8 lineage commitment. A developing αβ T cell commits to the CD4+ helper T cell fate or CD8+ cytotoxic T cell fate depending on the specificity of its TCR. A thymocyte expressing an MHC class I-restricted TCR will develop as a CD8+ T cell and one expressing an MHC class II-restricted TCR will develop as a CD4+ T cell. Classical models for lineage commitment posited that cells with higher TCR signaling intensity and/or longer TCR signal duration are directed toward the CD4+ lineage^66^. In contrast, the kinetic signaling model specifies that positively-selected DP thymocytes advance to a CD4+ CD8^lo^ selection intermediate^67,68^. If this intermediate continues to receive TCR stimulation despite the lack of CD8 expression, it differentiates into a CD4+ T cell. If the intermediate ceases to receive TCR stimulation, it becomes receptive to IL-7 signaling and undergoes co-receptor reversal (i.e., downregulation of *Cd4* and upregulation of *Cd8*). We observed that both CD4 SP and CD8 SP thymocytes were significantly reduced in the absence of NCOR1/2, although a 2-fold greater effect was observed for CD4 SP relative to CD8 SP thymocytes. As a result, the ratio of CD4+ to CD8+ T cells was significantly reduced in both the thymus and spleen. This finding indicates that NCOR1 and NCOR2 play a role in CD4/CD8 differentiation.

Increased or sustained TCR expression typically favors CD4 lineage commitment^69^. While we found evidence of increased TCR signaling in DKO thymocytes, we instead observed a bias toward the CD8 lineage. There are multiple potential explanations for the CD8 bias in DKO cells despite increased TCR signaling. First, the histone deacetylase HDAC3 has been shown to restrain CD8 lineage genes in DPs to allow for CD4 lineage commitment^70^. Because NCOR1 and NCOR2 are required to activate the deacetylase function of HDAC3, their absence may prevent HDAC3 from silencing CD8 lineage genes. Second, GATA3 promotes CD4 lineage commitment by inducing THPOK expression; THPOK in turn represses *Runx3* to inhibit CD8 lineage differentiation^71–73^. We noted increased expression of *Gata3* in the absence of NCOR1/2, which would otherwise be expected to promote CD4 differentiation and inhibit CD8 differentiation^74^. One possibility for this unexpected finding is that NCOR1/2 may interact with THPOK to repress *Runx3* transcription. While NCOR1 and NCOR2 have not been shown to interact with ZBTB7B (THPOK), they do interact with multiple ZBTB family members, including ZBTB7A, ZBTB16 (PLZF), and ZBTB27 (BCL6)^75–77^. However, a recent report from Bosselut and colleagues suggests that THPOK interacts with the NURD repressor complex and not NCORs^78^. Alternatively, GATA3 has been shown to repress *Runx3* independently of THPOK^79,80^. It is possible that GATA3 represses *Runx3*, and therefore CD4 differentiation^81,82^, by associating with NCORs: NCOR2 and GATA3 are predicted to interact^83^. In the absence of NCOR1/2, this GATA3-mediated repression of CD8 differentiation would no longer function. Third, expression of the TCR-induced transcription factor NUR77, which directly represses expression of *Runx3*^84^, was increased in DKO DP thymocytes. NUR77 is known to interact with NCOR2^84^. Without NCORs, NUR77 may no longer be able to repress *Runx3* expression, thereby promoting CD8 lineage commitment. We propose a model in which enhanced TCR signaling in the absence of NCOR1/2 results in increased GATA3 and NUR77 expression^42,74^. Although increased GATA3 and NUR77 expression would normally promote CD4 differentiation and inhibit CD8 differentiation, without NCOR1/2 corepressor complexes, they are unable to repress CD8 lineage commitment.

Loss of NCOR1/2 also affected rearrangement at the TCRα locus. Recombination events in the TCRα locus proceed in an “inside out” manner. Primary TCRα rearrangement occurs between the most proximal gene segments (3’ V_α_ and 5’ J_α_). The primary rearrangement directs increased chromatin accessibility to gene segments within several kilobases and decreases accessibility at further distances^85^. Secondary rearrangements use increasingly distal gene segments (progressively more 5’ for V_α_ segments and progressively more 3’ for J_α_ segments). Expression of RAGs continues in DP thymocytes until a cell dies or undergoes TCR signal-dependent positive or negative selection^46,47,86^. Using single cell TCR-seq, we identified a strong bias for proximal gene segments in NCOR-deficient thymocytes. We weighed multiple mechanistic explanations for the biased TCR repertoire. Because shortened DP thymocyte lifespan is associated with a proximal bias in the TCR repertoire, and NCOR-deficient thymocytes exhibit increased apoptosis, we examined the TCRα repertoire at the level of individual clusters^46,87^. In the DP-1 and DP-2 clusters, the usage of proximal TRAV and proximal TRAJ gene segments was not significantly different in the absence of NCORs. Premature death by neglect of DP thymocytes prior to distal rearrangements therefore is less likely to be the cause of the biased repertoire.

Decreased expression and/or function of RAGs following NCOR1/2 deletion might account for altered VJ recombination as this has been shown to promote a proximally-biased TCRα repertoire^88^. In the absence of NCORs, we found that *Rag2* expression was modestly decreased while *Rag1* expression was unchanged in NCOR-deficient thymocytes. Thus, reduced Rag2 expression may partially account for the reduced usage of distal V_α_ and J_α_ genes. Finally, structural changes in the absence of NCORs may contribute to the altered distribution of TCRα gene segments. TCRα rearrangement requires assembly of a chromatin loop comprised of CTCF, cohesin, *Tcra* enhancer (E_α_), and the T early α promoter (TEAp). CTCF and cohesin conditional KO mice do not recombine more distal TCRα segments^89,90^. NCORs may support the formation of the CTCF/cohesin complex. Supporting this conjecture, our scATAC-seq data from NCOR-deficient B cells demonstrated a major reduction in open chromatin for CTCF binding sites throughout the genome during B cell development^32^; a similar mechanism might operate in developing T cells as well. These data suggest multiple potential mechanisms for NCOR1/2 involvement in VJ rearrangement at the TCRa locus.

Our study also observed that NCOR1/2 collectively regulated thymocyte emigration. Importantly, all the genes needed to leave the thymus (including *Sell*, *S1pr1* and *Klf2*) were highly expressed in NCOR1/2 DKO CD4-SP2 thymocytes and the retained CD4SP subset. S1PR1 promoted lymphocyte egress from tissues while CD69 promotes their retention^38,50,54,55^. CD69 mediates internalization of S1PR1; co-expression of S1PR1 and CD69 on the cell surface should be mutually exclusive^54,57^. We were therefore surprised to observe co-expression of CD69 and S1PR1 on the surface of CD4 SP thymocytes from DKO mice. Our previous studies on B cells demonstrated that the *Cd69* gene locus is more accessible in the absence of NCOR1/2, which may account for the increased and prolonged CD69 expression. Why S1PR1 remained elevated in the presence of CD69 is less clear. Because the internalization of S1PR1 is a Gi-dependent process^54,57^, we considered which genes upregulated following Gi inhibition may be de-repressed in the absence of NCORs. The coupling between S1PR1 and Gi proteins is disrupted by pertussis toxin, blocking its inhibition of adenylyl cyclase^91^. This Gi inhibition results in the accumulation of cAMP and increased protein kinase A (PKA) signaling. PKA then regulates gene transcription by phosphorylating the transcription factors cyclic AMP-response element binding protein (CREB) and NF-κB^92,93^. NCOR1 and NCOR2 have been shown to interact with both of these PKA-activated transcription factors^9–11,94,95^. De-repression of CREB and/or NF-κB in the absence of NCOR1/2 may therefore yield a comparable phenotype to increased PKA activity (i.e., co-expression of CD69 and S1PR1). Through this mechanism, NCOR1/2 deletion may block the CD69-mediated internalization of S1PR1. Furthermore, CD69 acts as an agonist of S1PR1, rendering a cell unresponsive to an external S1P gradient and thereby blocking thymocyte egress^57^.

## Methods

### Animals

All mice in this study were bred and housed at the University of Minnesota under specific-pathogen-free conditions. Breeding and experimental protocols were approved by the Institutional Animal Care and Use Committee (2203-39908A; 2203-39912A). Animals were fed a standard rodent chow diet *ad libitum*. They were housed in a light/dark cycle of 14 hours/10 hours. Ambient temperature was maintained at 22°C, with humidity ranging from 30-40%. All transgenic mice were on the C57BL/6 background. *Ncor1^f/f^* mice were kindly provided by E. Olson (UT Southwestern, USA) and J. Auwerx (École Polytechnique Fédérale de Lausanne, Switzerland)^96^. A flip-excision strategy was used to generate *Ncor2* conditional knockout mice and was described in detail in Lee et al.^32,97^. *Cd4-cre* (022071), *Rosa26-LSL-tdTomato* (007914) and *Cd4-creER^T^*^2^ (022356) mice were purchased from the Jackson Laboratory^98–100^. All experiments in this study were performed with age-matched mice ranging from 6 to 10 weeks old.

### Tissue processing

Mice were euthanized using CO_2_ asphyxiation in accordance with our IACUC protocols. Thymi and spleens were harvested in FACS buffer (2% FBS with 2 mM ethylenediaminetetraacetic acid in PBS). Frosted glass slides were used to mechanically dissociate the tissue. Cells were filtered through a 70-um mesh filter and centrifuged at 350 g for 5 minutes. Thymocyte samples were resuspended in FACS buffer before proceeding with staining. Splenocytes were resuspended in ACK lysis buffer (0.15 M ammonium chloride, 10 mM potassium bicarbonate, and 0.1 mM EDTA), incubated at 22 C for 5 minutes, washed with FACS buffer, and then resuspended before proceeding with staining.

### Flow cytometry and antibodies

Single cell suspensions prepared from thymus and spleen samples were stained with the indicated antibodies. Antibodies used included: TCRβ (H57-597; 1:100), CD3 (17A2; 1:100), CD4 (GK1.5 and RM4-5; 1:100), CD5 (53-7.3; 1:100), CD8 (53-6.7; 1:100), CD25 (P61.5; 1:100), CD44 (IM7; 1:100), CD45 (30-F11; 1:100), CD69 (H1.2F3; 1:100), CD73 (TY/11.8; 1:100), I-A^b^ (AF6-88.5.5.3; 1:100), Sca-1 (D7; 1:100), Ly-6C (HK1.4; 1:100), Ly-6D (49-H4; 1:100), GITR (DTA-1; 1:100), S1PR1 (713412; 80 ug/mL), biotin mouse anti-rat IgG2a (RG7/1.30; 1:50), Foxp3 (FJK-16s; 1:100), NCOR1 (Cell Signaling Technology 5948; 1:100), donkey anti-rabbit IgG (eBioscience 12-4739-81; 1:100), BIM (C34C5; 1:50), and cleaved caspase-3 (D3E9; 1:50). For surface staining, cells were resuspended in FACS buffer containing antibodies at the dilutions specified above and incubated for 20 minutes at 4 °C. Cells were resuspended in FACS buffer and analyzed on a cytometer, or fixed and permeabilized for intracellular staining. For intracellular staining with Foxp3, NCOR1, and Bim antibodies, cells were resuspended in Fixation/Permeabilization working solution (eBioscience) and incubated at 22 C for 30 minutes. Cells were washed twice with Permeabilization Buffer (eBioscience) before resuspension in Permeabilization Buffer containing antibodies at the concentrations indicated above. Cells were stained for 30 minutes at 20 C. Intracellular staining with cleaved caspase-3 antibodies was performed as previously described^44^. In brief, after surface staining, cells were resuspended in Fixation/Permeabilization solution (BD Cytofix/Cytoperm kit) and incubated for 30 minutes at 4 C. After washing twice with BD Perm/Wash buffer, cells were stained with anti-cleaved caspase-3 for 30 minutes at 22 C. For S1PR1 surface staining, cells were stained with anti-S1PR1 antibody for 90 minutes on ice. After washing, cells were incubated with biotinylated mouse anti-rat IgG2a for 45 minutes on ice. Cells were again washed with FACS buffer and incubated with the remaining surface antibodies for 30 minutes at 4 C. Flow cytometry data were acquired using a BD LSR II or Fortessa cytometer (BD Biosciences) in the University of Minnesota flow cytometry core facility. All flow cytometry data were analyzed using FlowJo version 10.9 (FlowJo LLC).

### Thymic emigration assay

*Cd4-creER^T^*^2^ x *Ncor1^f/f^* x *Ncor2^f/f^* x *LSL-tdTomato* mice were administered 2 doses of tamoxifen by intraperitoneal injection (75 mg/kg), spaced 24 hours apart. Age- and sex-matched CD45.2 mice served as WT controls. Approximately 48 hours after the second dose of tamoxifen, intra-thymic biotin injections were performed using ultrasound guidance as previously described^58^. Briefly, mice were anesthetized using isoflurane gas and immobilized on a heated platform in the supine position. Depilatory cream was used to remove hair from the chest before applying ultrasound gel. A Vevo 2100 Imaging System (Visual Sonics) with a MS500 transducer was used to identify the thymus. The left and right thymic lobes were each injected with 20 uL NHS-biotin (1 mg/mL in PBS) under direct visualization. Mice were sacrificed two days later and thymi were processed for flow cytometric analysis as described above.

### Cell sorting and scRNA-seq

Thymi were removed from 6-week-old mice and mechanically dissociated using glass slides to prepare a single cell suspension. Cells were washed and resuspended in sort buffer (2% FBS, 5 mM EDTA in PBS). After reserving a small aliquot of each sample, we performed a CD8 depletion using anti-CD8α-FITC, anti-FITC MicroBeads (Miltenyi Biotec), and LS columns (Miltenyi Biotec). The reserved non-depleted cells and CD8-depleted samples were then stained with Viability Dye and a surface antibody cocktail diluted in sort buffer. The surface antibody cocktail included fluorescently-labeled antibodies for cell sorting as well as CITE-seq antibodies. CITE-seq antibodies included CD24 (TotalSeq-C0212), CD25 (TotalSeq-C0097), CD44 (TotalSeq-C0073), CD69 (TotalSeq-C0197), OX40 (TotalSeq-C0195), CCR4 (TotalSeq-C0833), CCR7 (TotalSeq-C0377), and TIGIT (TotalSeq-C0848). A unique hashtag antibody was used to label samples by mouse, which included TotalSeq-C0304 anti-mouse Hashtag 4, TotalSeq-C0305 anti-mouse Hashtag 5,TotalSeq-C0306 anti-mouse Hashtag 6, TotalSeq-C0308 anti-mouse Hashtag 8, TotalSeq-C0309 anti-mouse Hashtag 9, and TotalSeq-C0310 anti-mouse Hashtag 10 (all from BioLegend). 3 populations were sorted using a BD FACSAria II cell sorter (BD Biosciences). TCRβ+ CD4+ CD8+ (DP) cells were sorted from the non-depleted samples. TCRβ+ CD4+ CD8-CD73-CD25-GITR-cells (CD4 SP) and TCRβ+ CD4+ CD8-CD73-cells that were CD25+ and/or GITR+ (Tregs and Treg progenitors) were sorted from the CD8-depleted samples. These populations were collected in equal proportion for each mouse. Cells were sorted into 50% FBS in 1x PBS. Samples were pooled, washed with capture buffer (10% FBS in PBS), and resuspended in capture buffer prior to library preparation.

### Single-cell RNA and TCR sequencing

For scRNA-seq, we generated three 5’ libraries that measure (1) mRNA expression (RNA), (2) feature barcodes, including hashtag and CITE-seq antibodies (FB), and (3) TCR sequencing (TCR). The prepared sample was split into three libraries (RNA, FB, and TCR). Reverse transcription PCR and library preparation were carried out under the Chromium Single Cell 5’ v2 protocol (10X Genomics) according to manufacturer specifications. After library preparation, quality control was performed using a bioanalyzer (Agilent) and MiSeq low-depth sequencing (Illumina) to determine the approximate number of cells and estimate quality of the dataset. The library was sequenced on a NovaSeq 6000 (Illumina) with 2 x 150-bp paired-end reads. Sequencing depth was 50,000 reads per cell. Raw and processed data have been deposited at Gene Expression Omnibus.

### Single-cell bioinformatic analysis

The single cell RNA transcriptomic (GEX) sequencing data was processed using the CellRanger pipeline version 6.0.0 (10X Genomics). Reads were demultiplexed using ‘cellranger mkfastq’ and aligned to the mouse genome (mm10, refdata-gex-mm10-2020-A, provided by 10X Genomics) using ‘cellranger count’. Raw counts were analyzed using the Seurat R package (v 4.1.0). First, the data set was filtered to include GEMs (gel beads in emulsion) that contained 1 cell using the Seurat function ‘HTODemux’. A k-value of 7 showed the cleanest results (11,216 cells). Second, the filtered RNA data was log normalized and scaled based on the top 2000 most variable genes. Principle component analysis (PCA) was performed on the normalized and scaled data, and the top 30 PCA vectors were used as input to create a UMAP (‘RunUMAP’). Cells were then clustered using the ‘FindNeighbors’ and ‘FindClusters’ functions. A range of possible resolutions were tested, ranging from 0.1 to 1.5. The ‘BuildClusterTree’ function was used to aid in selecting resolutions for differential expression (DE) analysis using the FindMarkers function. Both individual cluster to all clusters and pairwise cluster-to-cluster comparisons were used for DE analysis. Final cluster classification for the mixed (WT, NCOR1 KO, DKO) genotype population was based on resolution 1.4.

### Single-cell T cell receptor sequencing analysis

The single cell T-cell VDJ (TCR) sequencing data was processed using the CellRanger pipeline (10X Genomics, v. 6.0.0). Reads were demultiplexed using ‘cellranger mkfastq’ and reads were mapped to a V(D)J reference (provided by 10X Genomics, mm10 refdata-cellranger-vdj-GRCm38-alts-ensembl-5.0.0) and assembled into nucleotide-based clonotypes using ‘cellranger vdj’. Custom R code (v 4.0.4) was used to filter and collect J and V gene (both alpha and beta gene) sequences, merged with the metadata from the mixed genotype (WT, NCOR1, NCOR-DKO) single cell gene expression data (resolution 1.4 clusters) and then tallied for genotype and cluster and ordered by gene coordinates retrieved from the mm10 genome (v. Mus_musculus.GRCm39.106). The code used for this study is available upon request.

### Gene set enrichment analysis

Ranked DE gene lists were generated using the Seurat ‘FindMarkers’ function. The R-based program ClusterProfiler (4.8.2) was used to perform gene set enrichment analysis (GSEA) to identify gene sets from the Molecular Signature Database (MSigDB, v 7.5.1) that are enriched for single cell DE lists^101,102^.

### Statistical analysis

All statistical analysis with flow cytometry-derived data was conducted using GraphPad Prism v9.5.1 (GraphPad Software, USA), with the corresponding statistical tests and multiple comparison corrections listed in the figure legends. Differential gene expression analysis was completed using the FindMarkers function. Differential expression was determined to be significant if the adjusted p value was 0.05 or lower and the log2FC was at least 0.2.

## Acknowledgements

We thank G. Hubbard, T. Maiers, M. Dinh, and L. Heltemes-Harris for their technical assistance with mouse husbandry and colony management; E. Stanley and K. Beckman from the University of Minnesota Genomics Center for single-cell capture and sequencing; T. Knutson at the Minnesota Supercomputing Institute; J. Motl and P. Champoux from the University Flow Cytometry Resource; J. Mitchell from University Imaging Centers. This work was supported by an NIH individual predoctoral F31 fellowship (F31 AI157166) and a T32 training grant (T32 AI007313) to N.A.D., predoctoral fellowship (F30 CA232399) and training grant (T32 GM008244) to R.D.L, and NIH grant R01 AI147540 to M.A.F.

---

**Supplemental Figure 1.**
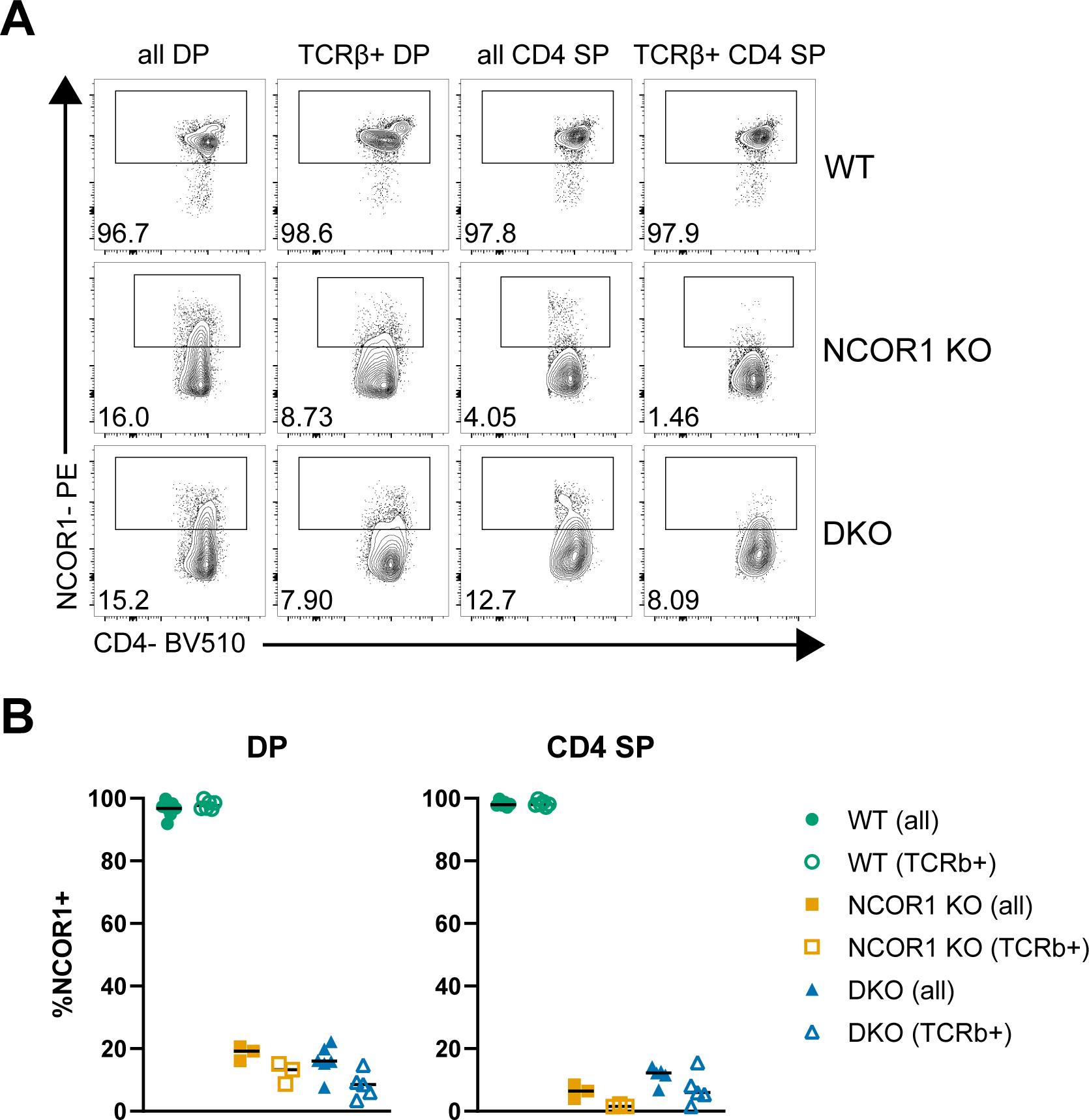
Efficiency of NCOR1 deletion in DP and CD4 SP thymocytes. (A) Representative flow cytometry plots demonstrating residual NCOR1 expression in thymocytes isolated from wildtype (WT; top row), *Cd4-cre x Ncor1^f/f^* (NCOR1 KO; middle row), and *Cd4-cre x Ncor1^f/f^ x Ncor2^f/f^* (DKO; bottom row) mice. Pre-gated on live CD4+ CD8+ cells (left panel), live TCRβ+ CD4+ CD8+ (second from left), live CD4+ CD8-(second from right), and live TCRβ+ CD4+ CD8-(right panel) cells. (B) Summary data of NCOR1+ frequency for populations as described in A. Data were compiled from 4 independent experiments. Horizontal bars represent the median.

**Supplemental Figure 2.**
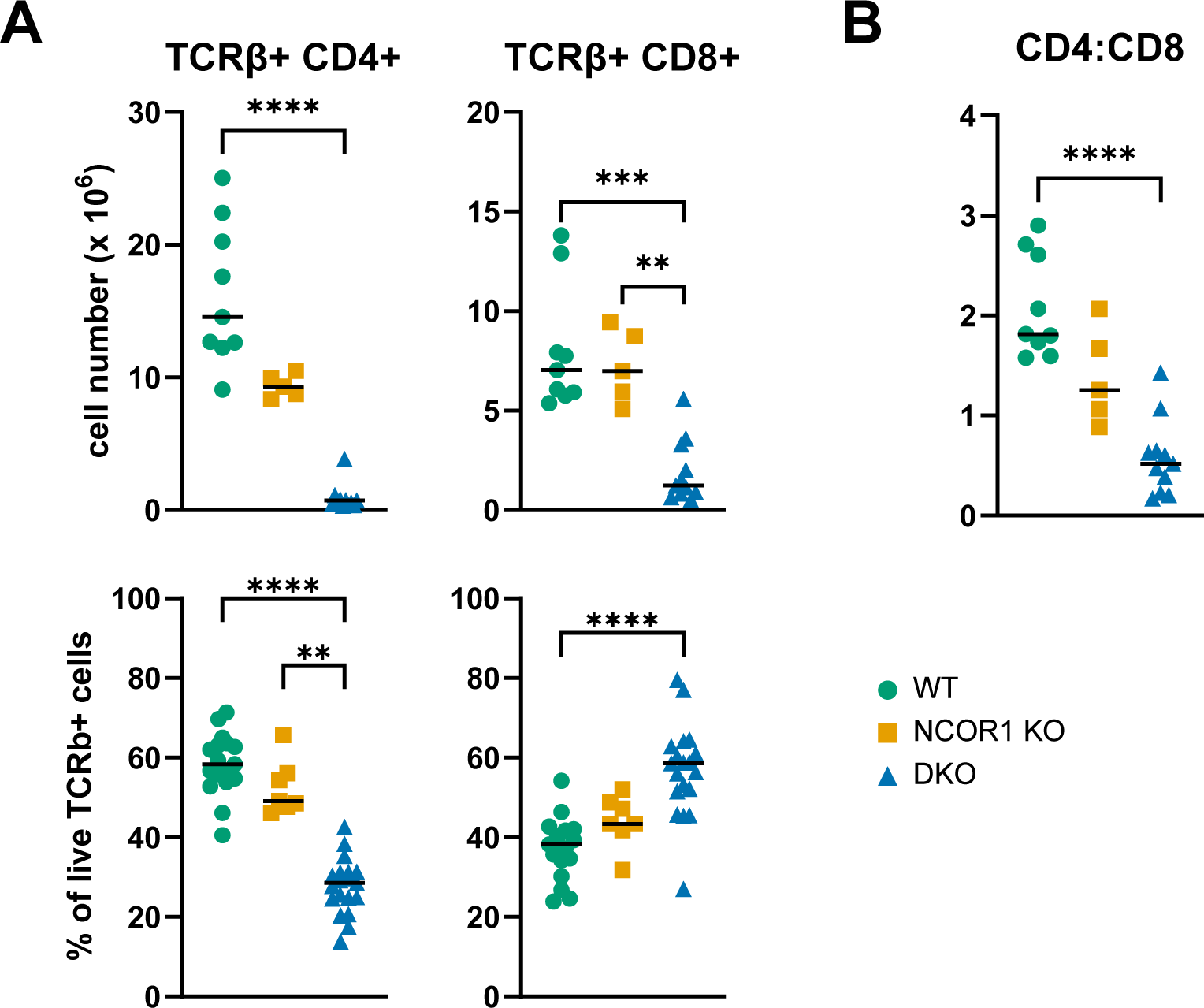
Impact of loss of NCOR1 and NCOR2 in the spleen. (A) Summary data of TCRβ+ CD4 and TCRβ+ CD8 cell counts (top panel) and frequencies (bottom panel) in splenocytes isolated from WT, NCOR1 KO, and DKO mice. (B) Ratio of TCRβ+ CD4 T cells to TCRβ+ CD8 T cells in the spleen. Data were compiled from more than 5 independent experiments. Horizontal bars represent the median. Kruskal-Wallis test with Dunn’s multiple comparison test was used to assess statistical significance. *p < 0.05, **p < 0.01, ***p < 0.001, ****p < 0.0001.

**Supplemental Figure 3.**
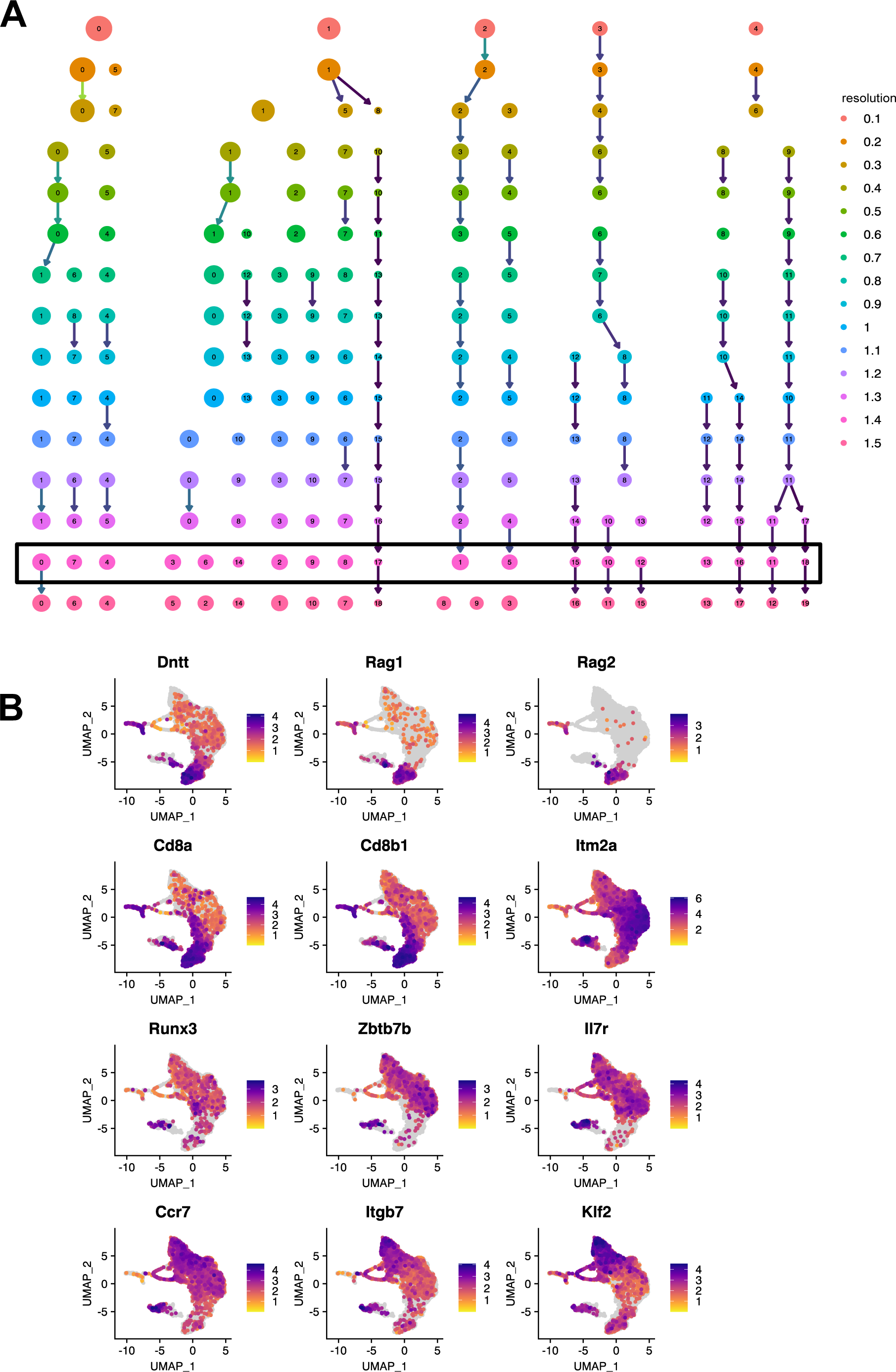
scRNA-seq cluster resolution comparison and expression patterns for genes of interest. (A) Clustree plot highlighting the selected resolution of 1.4 for analysis of full dataset (all genotypes combined). (B) scRNA-seq feature plots of genes involved in TCR recombination (*Dntt*, *Rag1*, *Rag2*), DP (*Cd8a*, *Cd8b1*), lineage specification (*Runx3*, *Zbtb7b*), positive selection (*Ccr7, Itm2a, Il7r*), and maturation (*Itgb7, Klf2*).

**Supplemental Figure 4.**
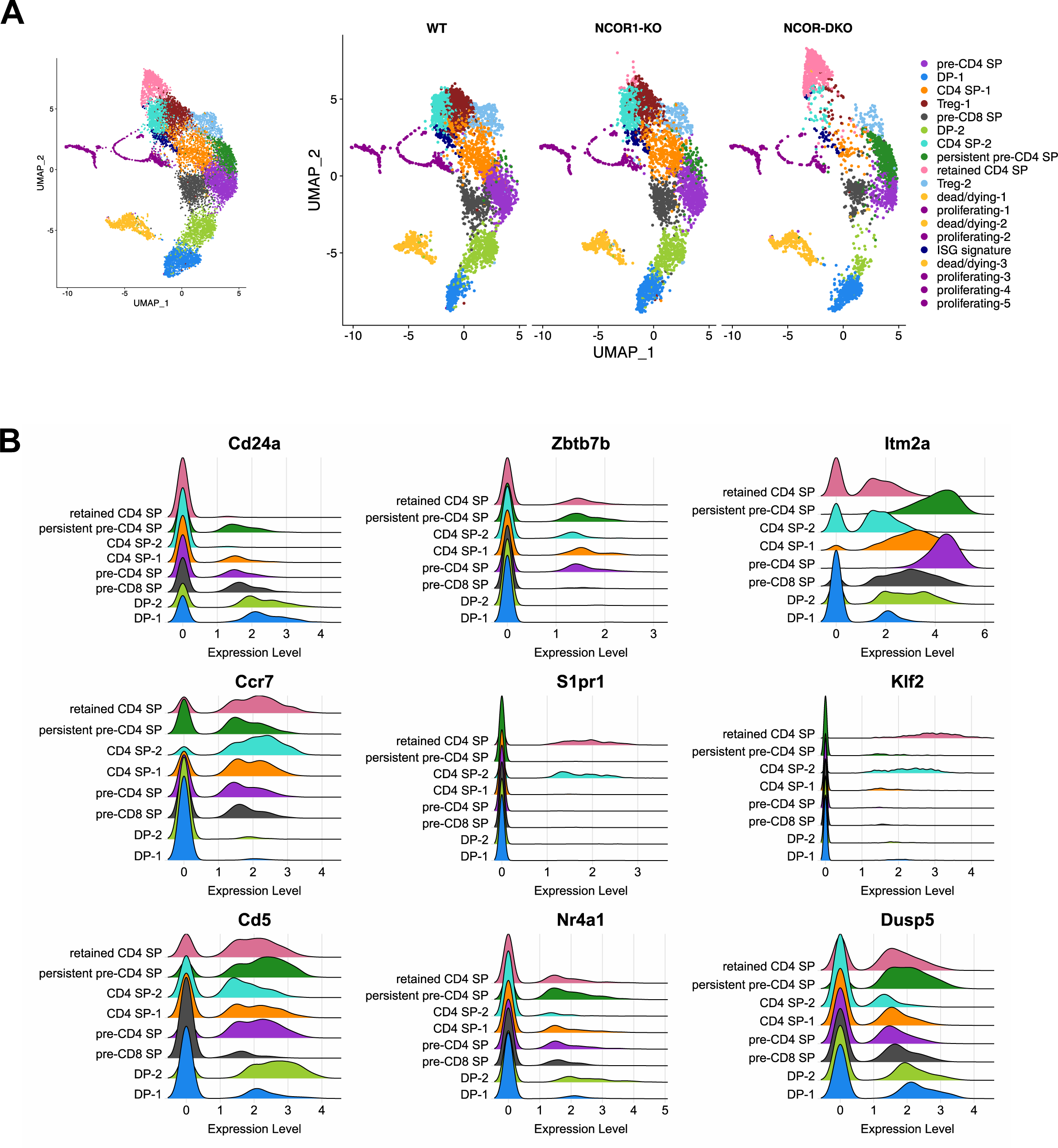
scRNA-seq cluster identification. (A) UMAP plot for combined dataset (left) and UMAP plots separated by genotype (right) with full list of cluster identities. (B) Ridge plots for select genes related to maturity, selection, and TCR signaling to compare expression patterns of DKO-predominant clusters (persistent pre-CD4 SP, retained CD4 SP) to clusters with more proportionate representation of WT cells (DP-1, DP-2, pre-CD8 SP, pre-CD4 SP, CD4 SP-1, CD4 SP-2).

**Supplemental Figure 5.**
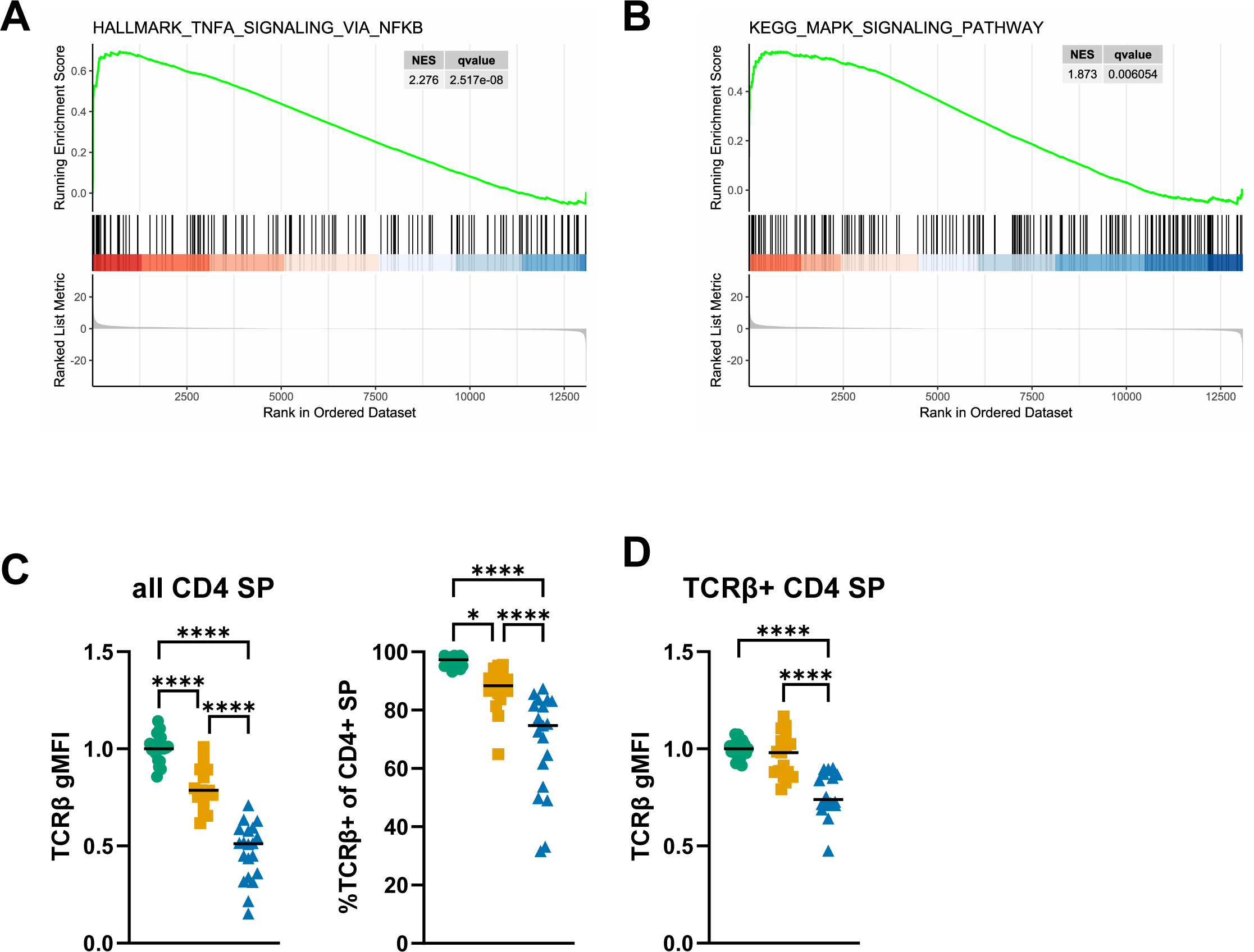
Changes in cell signaling pathways and TCRβ expression in the absence of NCOR1/2. (A) GSEA for hallmark TNFα signaling via NF-κB gene set in the pre-CD4 SP cluster. (B) GSEA for Kegg MAPK signaling in the pre-CD4 SP cluster. (C) Compiled gMFI data for TCRβ expression in all CD4+ SP thymocytes (left). Proportion of TCRβ+ cells among all CD4 SP thymocytes (right). (D) Compiled gMFI data for TCRβ expression in TCRβ+ CD4+ SP thymocytes. All gMFI values were normalized to WT sample(s) within each experiment. Horizontal bars indicate median. Flow cytometry data were compiled from more than 5 independent experiments. Statistical significance was determined by one-way ANOVA with Tukey’s multiple comparison test. *p < 0.05, **p < 0.01, ***p < 0.001, ****p < 0.0001.

**Supplemental Figure 6.**
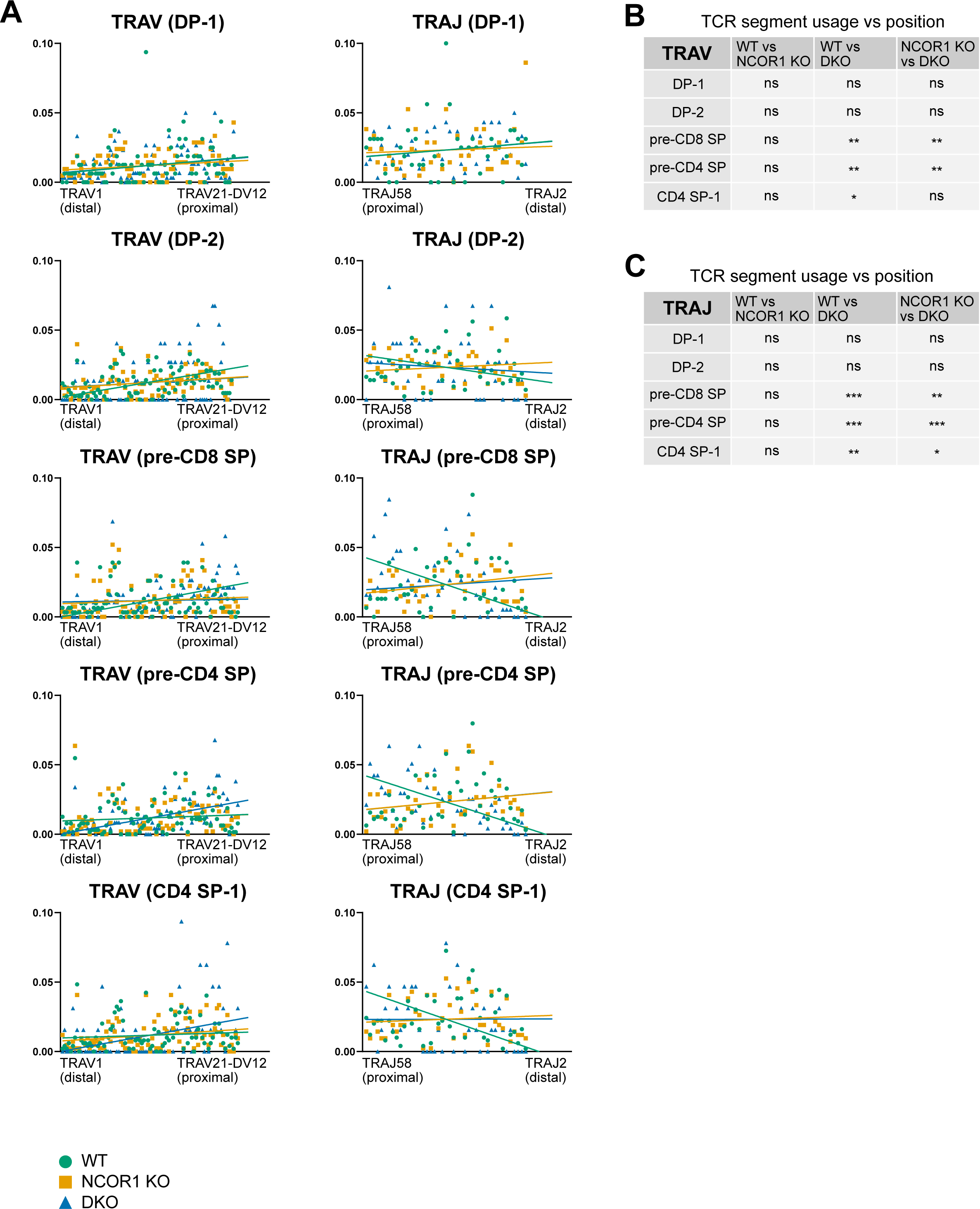
Altered TCRα repertoire is not driven by DP thymocytes. (A) Linear regressions of TRAV (left column) and TRAJ (right column) segment position, relative to initial recombination site, and proportion of cells with this segment found in each genotype. Each plot is labeled with the name of the cluster. (B) Comparison of relationship between distribution of TRAV segment utilization and position (slope from linear regressions as in panel A across individual clusters. (C) Comparison of relationship between distribution of TRAJ segment utilization and position across individual clusters. Statistical significance was determined by one-way ANOVA with Tukey’s multiple comparison test using slopes from linear regression (B, C). *p < 0.05, **p < 0.01, ***p < 0.001.

**Supplemental Figure 7.**
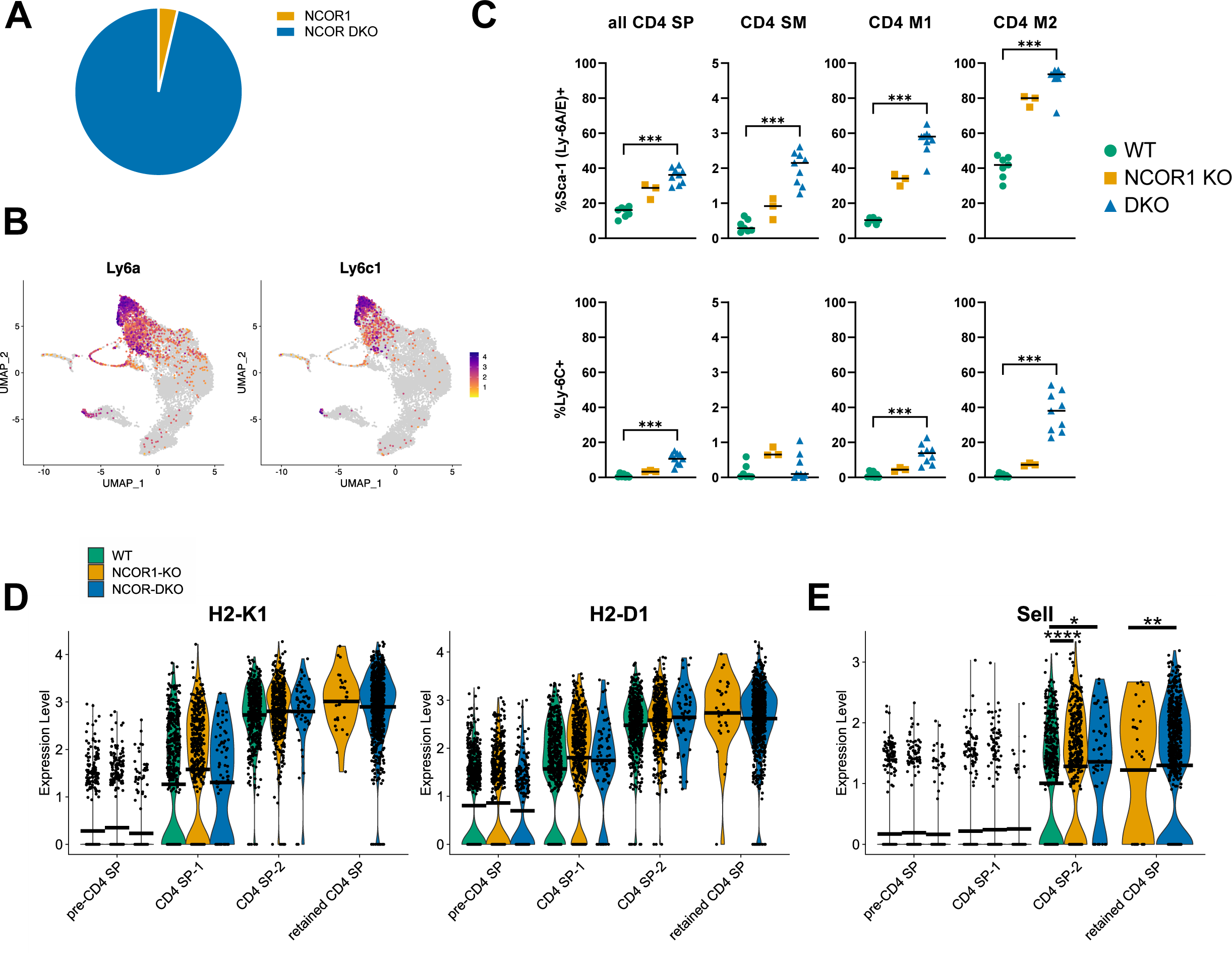
Retained CD4 SP. (A) Genotype distribution of cells within the retained CD4 SP cluster (0 for WT, 29 for NCOR1 KO, and 763 for DKO). (B) Feature plots for the top differentially upregulated genes in this cluster of the scRNA-seq dataset: *Ly6a* (left) and *Ly6c1* (right). (C) Ly-6A+ (top) and Ly-6C+ (bottom) expression in cells by flow cytometry in all TCRβ+ CD4 SP (left), CD4 SM (second from left), CD4 M1 (second from right), and CD4 M2 (right) thymocytes. Gating for CD4 SP subpopulations as in Figure 1D. (D) Violin plot demonstrating changes in *H2-K1* (left) and *H2-D1* (right) expression as CD4 SP thymocytes mature, compared to expression in retained CD4 SP population. (E) Violin plot demonstrating changes in *Sell* (CD62L) expression over the course of CD4 SP thymocyte maturation. Horizontal bars indicate median (C) or mean (D,E). Flow cytometry data was compiled from 5 independent experiments. Statistical significance was determined by Kruskal Wallis test with Dunn’s multiple comparison test (C). *p < 0.05, **p < 0.01, ***p < 0.001, ****p < 0.0001.

